# A pathogenic fungus uses volatiles to entice male flies into fatal matings with infected female cadavers

**DOI:** 10.1101/2021.10.21.465334

**Authors:** Andreas Naundrup, Björn Bohman, Charles A. Kwadha, Annette B. Jensen, Paul G. Becher, Henrik H. De Fine Licht

## Abstract

To ensure dispersal, many parasites and pathogens behaviourally manipulate infected hosts. Other pathogens and certain insect-pollinated flowers use sexual mimicry and release deceptive mating signals. However, it is unusual for pathogens to rely on both behavioural host manipulation and sexual mimicry. Here, we show that the host-specific and behaviourally manipulating pathogenic fungus, *Entomophthora muscae*, generates a chemical blend of volatile sesquiterpenes and alters the level of natural host cuticular hydrocarbons in dead infected female house fly (*Musca domestica*) cadavers. Healthy male house flies respond to the fungal compounds and are enticed into mating with dead female cadavers. This is advantageous for the fungus as close proximity between host individuals leads to an increased probability of infection. The fungus-emitted volatiles thus represent the evolution of an extended phenotypic trait that exploit male flies’ willingness to mate and benefit the fungus by altering the behavioural phenotype of uninfected healthy male host flies.

## Main

The evolution of specific mate recognition systems is often central for successful sexual reproduction (Ryan and Rand, 1993). Once males and females have located each other, individual mating preferences or competition for access to mates may lead to suboptimal decisions during courtship and mating (Trivers, 1972; Andersson, 1994). The willingness to mate is for example exploited by certain insect pollinated flowers (Schiestl *et al*., 2000; Cohen *et al*., 2021; Hayashi *et al*., 2021), which use sexual mimicry to attract pollinators by resembling the opposite sex visually and/or chemically. Exploitation of mate recognition systems can be highly advantageous for obligate pathogens as it increases the chance of pathogen transmission by ensuring contact with new potential hosts of the right species (Hansen and De Fine Licht, 2019).

Obligate parasites and pathogens are under strong selection pressure to find a host to continue or complete their life-cycle (Schmid-Hempel, 2011). This has led to the convergent evolution of behavioural manipulation across parasitic phyla that increases transmission to new hosts (Helluy and Thomas, 2003; Hoover *et al*., 2011; Adamo, 2013; de Bekker *et al*., 2015; Ros *et al*., 2015; Botnevik *et al*., 2016; Małagocka, Jensen and Eilenberg, 2017). Pathogens may also behaviourally manipulate their hosts without residing inside the body of the host (Hughes and Libersat, 2018), for example through host-consumption of pathogen secreted substances (Hojo, Pierce and Tsuji, 2015) or via direct injection of chemical cocktails by certain parasitoids (Gal and Libersat, 2010). A widespread, but more subtle behavioural manipulation is attraction of potential new hosts or vectors with volatile compounds by certain entomopathogenic bacteria (Keesey *et al*., 2017), nematodes (Zhang *et al*., 2019), fungi (George *et al*., 2013; Trandem *et al*., 2015), beetle-tapeworms (Evans *et al*., 1998; Shostak and Smyth, 1998; Shea, 2007), and plant viruses (Mauck, De Moraes and Mescher, 2010). Such attraction is driven by the extended phenotype of pathogens in the infected host (Dawkins, 1982; Hoover *et al*., 2011), where expression of pathogen genes ultimately leads to the altered behaviour of uninfected hosts from a distance. Although the proximate mechanism is in many cases still largely elusive (van Houte, Ros and van Oers, 2013; Hughes and Libersat, 2018; de Bekker, Beckerson and Elya, 2021), a combination of pathogenic traits governing the extended phenotype and the exploitation of host compensatory responses to pathogen infection is thought to control altered host behaviour (Lefèvre *et al*., 2009).

The entomopathogenic fungus, *Entomophthora muscae*, represents a unique example in that it not only behaviourally manipulates its immediate house fly (*Musca domestica*) host, but also appears to manipulate uninfected conspecifics. Prior to death, infected house flies are manipulated to seek out elevated positions and die with wings spread out in a specific posture conducive for active discharge of infectious fungal conidia, which occurs primarily within the first 24 hours post mortem (De Ruiter *et al*., 2019; Lovett *et al*., 2020). Male and female flies in the vicinity may become infected from being directly hit by a conidium (spore) or via contact with already-discharged conidia that form a halo on the surface surrounding a conidia-shooting cadaver (De Ruiter *et al*., 2019).

Remarkably, the behavioural manipulation by *E. muscae* continues after death of female hosts, as the fungus appears to use sexual mimicry to lure healthy males to attempt mating with these cadavers (Møller, 1993), even though they are only rewarded with deadly fungal conidia for their efforts (Fig. 1A). In house flies, courtship starts with the male jumping on top of the female in a so-called “mating strike” and placing his legs at the base of the wings of the female, which in turn instantly spreads her wings horizontally from the body resembling flight position (Murvosh, Fye and LaBrecque, 1964; Tobin and Stoffolano, 1973). Both males and females are considered as choosing sex in house flies because males vary considerably in their mating efforts and females can exert mating choice by kicking off courting males (Goulson *et al*., 1999). Although house fly males are able to distinguish between male and female cadavers covered in infectious *E. muscae* conidia (Zurek *et al*., 2002), males do not discriminate the fungus-swollen female cadavers and appear to readily initiate courtship and mating strikes (Møller, 1993).

**Fig. 1.**
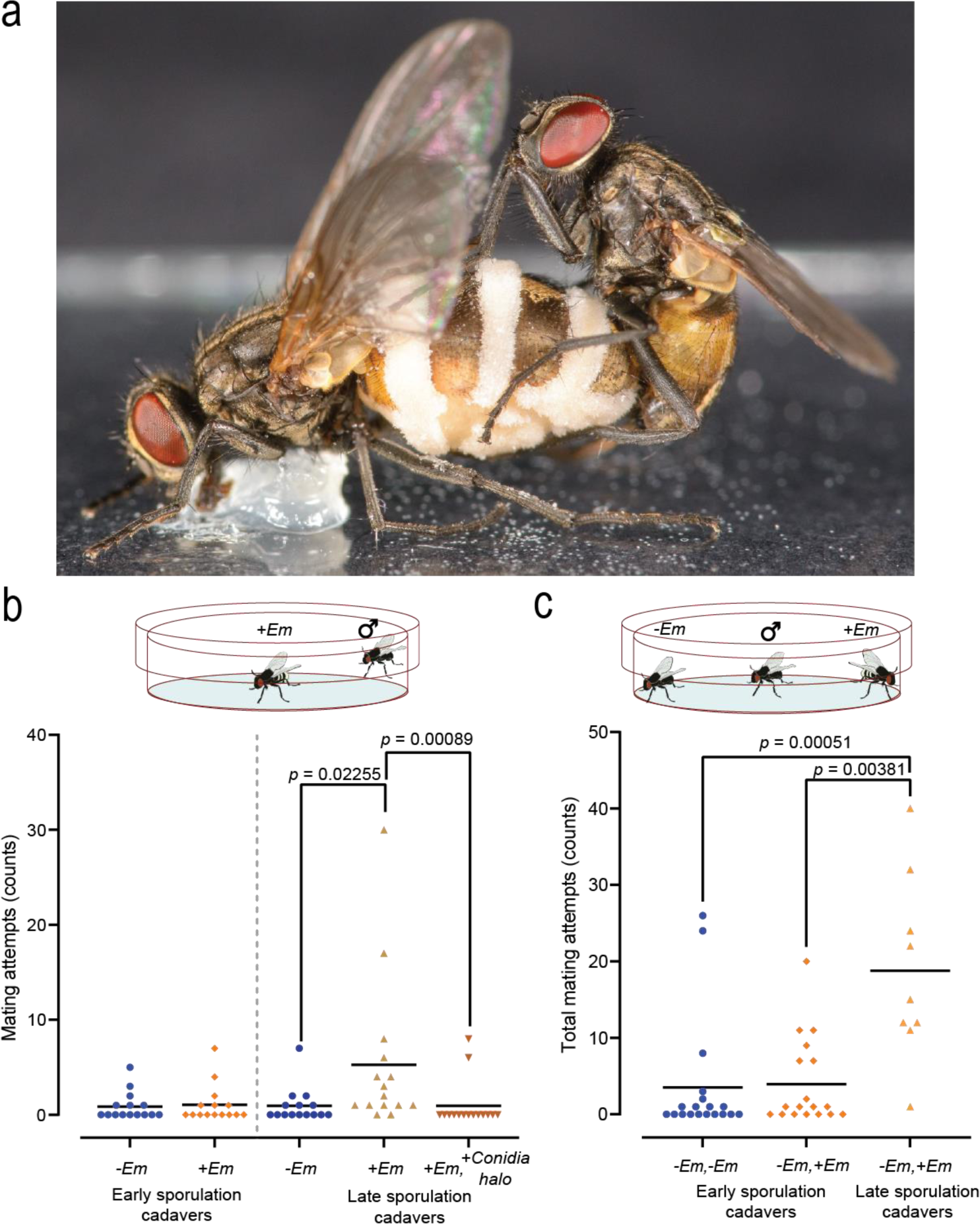
Male house fly mating attempts towards *E. muscae* cadavers. **a**, Healthy male house fly attempting to mate with *E. muscae* sporulating cadaver. Fungal growth is seen as white bands (conidiophores with conidia) extruding from the abdomen of the dead female. The actively discharged conidia are covering large parts of the wings and body of the female cadaver and also create a halo of conidia around the cadaver (Photo: Filippo Castelucci). **b**, Male mating attempts towards uninfected freeze-killed (*-Em*) or infected (*+Em*) *E. muscae*-killed cadavers in early (3-8 hours post death) and late (26-28 hours post death) sporulation stages (n = 15 per treatment). **c**, Total mating attempts by a male towards either cadaver when given a choice between two female cadavers that both were either uninfected (*-Em, –Em*), one uninfected and one infected in early sporulation stage (*-Em, +Em*), or one uninfected and one infected in late sporulation stage (*-Em, +Em*) (n = 9-19 per treatment).

We sought to understand the mechanisms governing this maladaptive behaviour, and test the hypothesis that an extended phenotype of *E. muscae* lures male flies by changing the chemistry of infected female cadavers. Specifically, we examined two fungal attraction pathways: 1*) E. muscae* amplifies the signal of female house fly pheromones, which creates a sensory bias so males respond to a quantitatively improved supernormal stimuli (amplified chemical attraction), or 2) *E. muscae* synthesizes new chemical cues not normally occurring in female house flies so males respond to a qualitatively improved mating signal (novel chemical attraction). Mate recognition in house flies is normally governed by both visual and chemical cues (Rogoff *et al*., 1964; Adams and Holt, 1987), and sex pheromones consist of sex-specific blends of cuticular hydrocarbons (CHC) that line the outer cuticle (Carlson *et al*., 1971; Adams, Nelson and Fatland, 1995; Noorman and Otter, 2001). Most CHCs are only perceived at close range (Blomquist, Tittiger and Jurenka, 2020), however, chemical derivatization can increase volatility and result in attraction over longer distances as shown in fruit fly (*Drosophila melanogaster*) pheromones (Lebreton *et al*., 2017).

We evaluated the potential amplified or novel chemical attraction pathways in three steps. First, we quantified male sexual attraction to fungus-killed cadavers and fungal conidia using behavioural assays. Second, we identified the chemical cues eliciting male mating attraction using chemical analyses (GC-MS) and physiological mechanisms enabling males to detect these cues using electroantennography (GC-EAD). Third, we verified the fungus *E. muscae* as source of the behaviourally active volatile compounds in fungus-killed cadavers using transcriptional profiling (RNAseq) of expressed genes in volatile chemical biosynthesis pathways.

## Results

### Male houseflies attempt mating with fungus-killed cadavers and are attracted to *E. muscae* conidia

We first performed mating activity experiments to verify increased male mating attempts towards *E. muscae*-killed female cadavers (Møller, 1993; Zurek *et al*., 2002) (Fig. 1A, suppl. Fig. 1, 2). Males neither spent more time in the vicinity of, nor physically touched fungus-killed cadavers more often than control cadavers (Supp. Fig. 3). This was irrespective of whether the cadavers were in early (3-8 hours post mortem) or late (26-28 hours post mortem) sporulation stages (Supp. Fig. 3). However, male mating attempts increased as the cadavers proceeded to the late stage of sporulation and were not surrounded by a halo of infective conidia (Kruskal-Wallis (K-W), χ^2^ = 14.095, p = 0.0008695, Fig. 1B). This increase in sexual attraction appeared sex-specific; males mated more frequently only with female cadavers (Suppl. Fig. 3, 4), confirming previous observations (Zurek *et al*., 2002). Interestingly, when we allowed a male to choose between a control and infected female cadaver present together in a small arena, we did not see a significant difference between infected and uninfected cadavers in the number of mating attempts (Suppl. Fig. 5). However, a significantly higher number of total mating attempts to either of the two cadavers in the arena was observed if just one of the two cadavers was in the late stage of sporulation (K-W χ^2^ =15.007, p = 0.0005512, Fig. 1C), which indicates that although the male did not discriminate between infected or uninfected female cadavers, the presence of mating cues in late-stage sporulating female cadavers stimulate male sexual behaviour. To confirm that manipulation of male sexual behaviour is an effective means of increasing fungal transmission, the male subjects were incubated for 10 days, which showed that 73% become infected after exposure to late sporulation cadavers compared to 15% of males exposed to early sporulation cadavers (Suppl. Data 1). Increased cadaver mating attempts correlated with fungal infection as more male flies became infected when exposed to a late sporulation stage female cadaver compared to a late sporulation stage male cadaver (Suppl. Data 1).

**Fig. 2.**
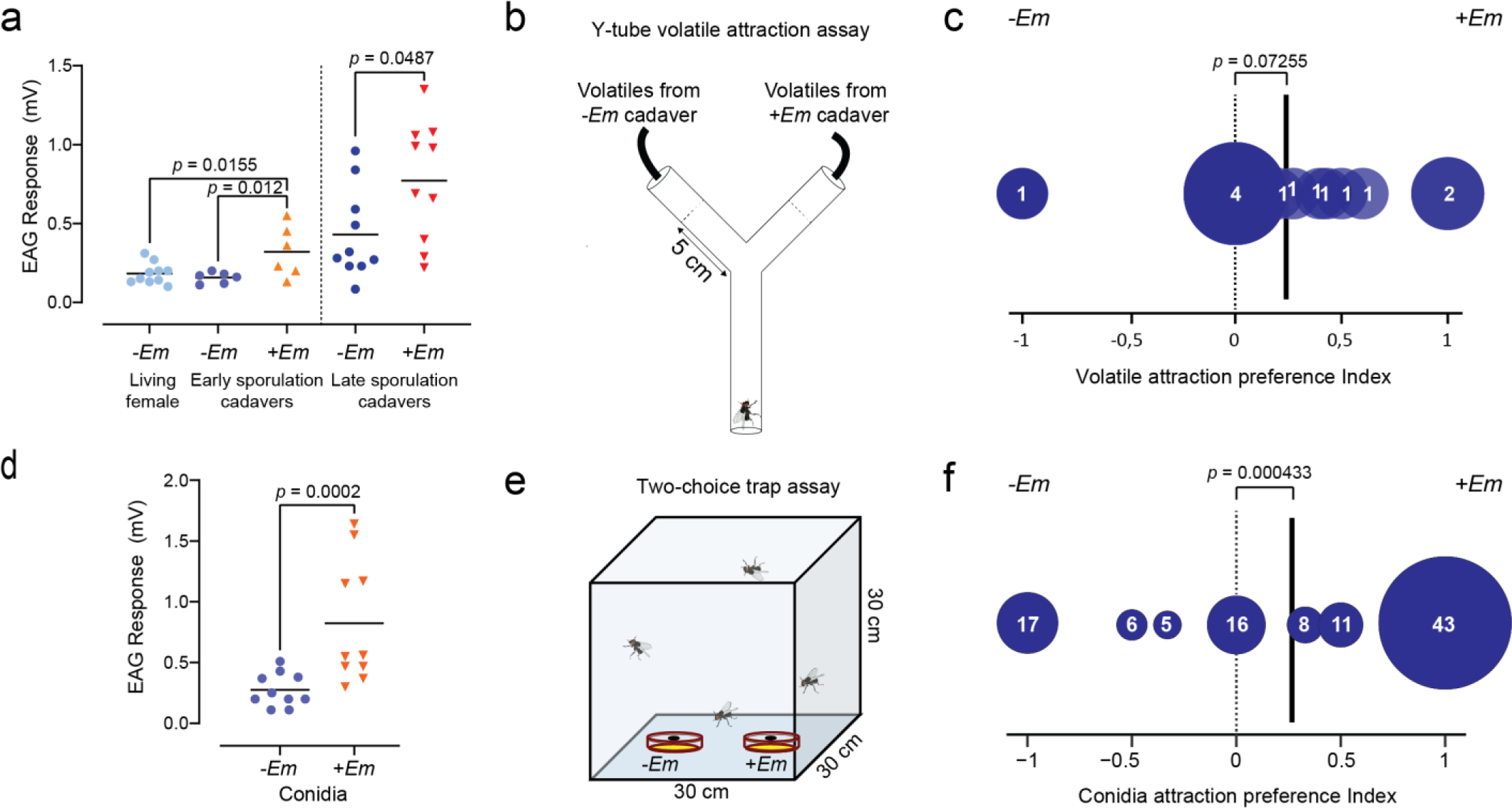
Antennal and behavioural responses to *E. muscae* volatile compounds. **a**, Male antennal EAG responses (mV) to volatile chemical blends of uninfected (-*Em*) or infected (+Em*) E. muscae-*killed cadavers in early (3-8 hours post death) and late (26-28 hours post death) sporulation stages (n = 6-10 per treatment). **b**, Drawing of Y-tube assay set-up. When the test fly reached 5 cm into either arm a choice was noted. **c.** Preference index of male housefly attraction in Y-tube experiments to volatile blends from *E. muscae* infected female cadavers (+*Em*) vs. volatiles from uninfected control female cadavers (*-Em*). Size of circles and numbers within show number of one-hour trials resulting in a given preference index (n = 3-12 male flies per trial, in total n = 90 flies). Solid black vertical lines designate mean preference index. **d**, Male antennal EAG responses (mV) to volatile chemical blends from *E. muscae* conidia and control (n = 10). **e**. Drawing of conidia attraction assay. **f.** Preference index of housefly attraction to volatile chemicals being emitted from visually obstructed sticky-traps with (*+Em*) and without *E. muscae* conidia (*-Em*). Size of circles and numbers within show number of 24-hour trials resulting in a given preference index (n = 4 male or female houseflies per trial, in total n = 106 trials / 424 flies tested).

**Fig. 3.**
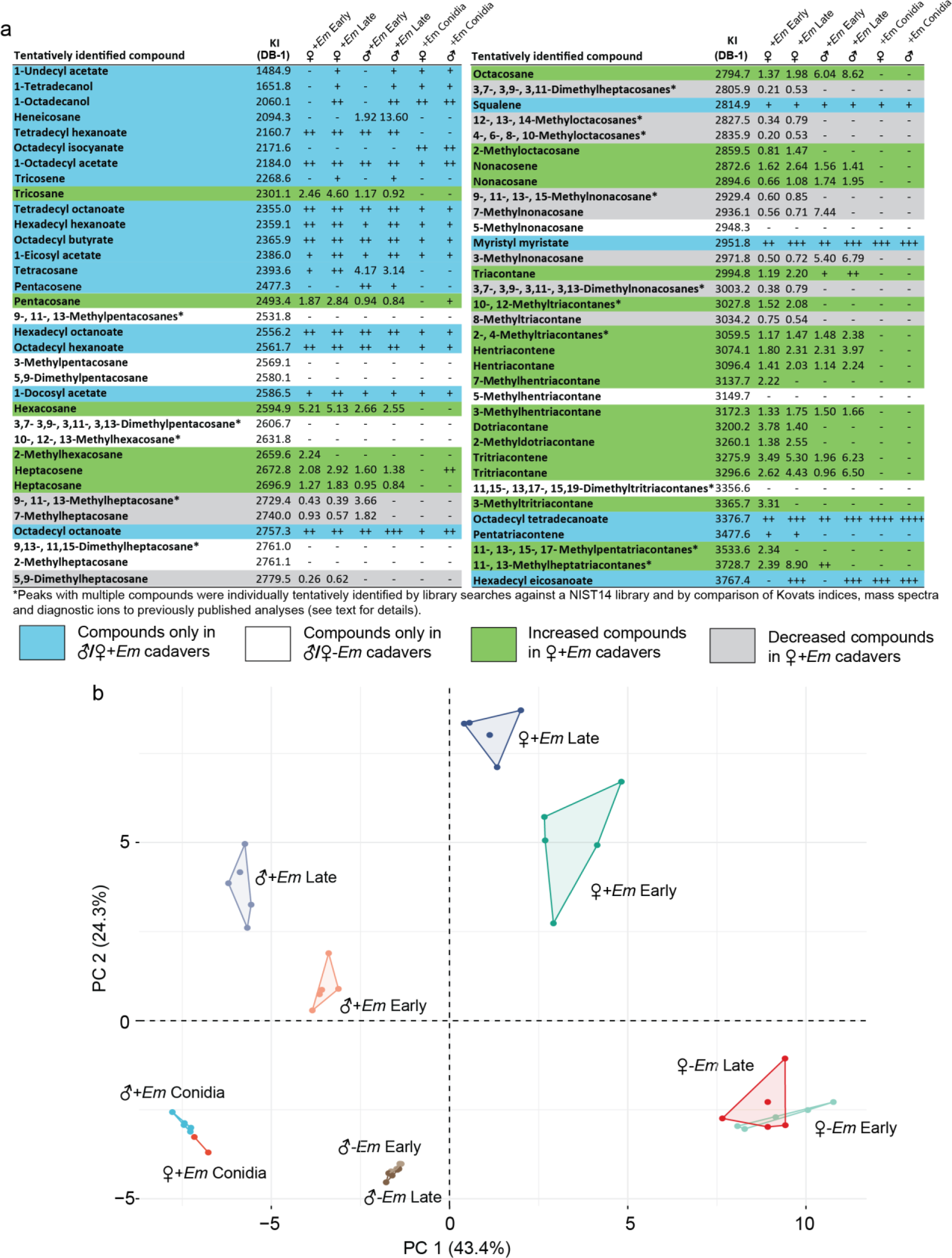
Cuticular chemical profile of *E. muscae* sporulating house flies. **a**, Tentatively identified compounds in cuticular hexane extracts of early (3-8 hours post death) and late (26-28 hours post death) female and male sporulating cadavers and conidia. Numbers denote fold change in intensity of total ion chromatogram (TIC) compared to corresponding uninfected controls, whereas + denotes presence in the sample, but not in corresponding control (+: < 5.5x107, ++: 5.51x107 < 5.5x108, +++: 5.51x108 < 5.5x109, ++++ > 5.51x109), and – denotes absence (i.e. only found in uninfected control samples). Compounds are colour coded whether they increase (green) or decrease (grey) in female *+Em* cadavers, or exclusively are found in *+Em* (blue) or *–Em* (white) cadavers. For each compound the retention index (KI, DB-1 column) is given. **b**, Principal component analysis (PCA) of cuticular and conidia extracts (hexane) shown in **a**.

During our behavioural studies, we occasionally observed males being attracted to conidia and approach these on artificial surfaces even in the absence of a cadaver. Here, the males would extend their proboscis and taste *E. muscae* conidia (Suppl. Figure 1). To investigate whether conidia alone were sufficient for attraction, flies were allowed to choose between two visually obstructed sticky traps inside a cage (Fig 2E). One trap contained *E. muscae* conidia from three all-male or all-female cadavers and the other was a control trap without conidia (exposed to uninfected cadavers). Both male and female flies were caught most often in traps with *E. muscae* conidia (logistic regression, n = 117, Z = 3.519, p = 0.000433, Fig. 2F), regardless of the sex of the conidia-discharging cadavers. As the traps were visually obstructed, we hypothesized that the attraction towards conidia was volatile-mediated and we therefore used electroantennography (EAG) to measure the male house fly antennal response to volatile compounds surrounding living flies, uninfected cadavers, sporulating cadavers, or conidia (headspace sampling) (Fig. 2A,D). Male house flies showed significantly higher antennal responses to conidial headspace (Linear mixed effects model (LMM), t = 15.728, p < 0.0001, Fig. 2D), and *E. muscae* sporulating cadaver headspace compared to headspace from either live (LMM, t = 3.573, p = 0.0112, Fig. 2A) or dead control flies (LMM, t = 3.914, p = 0.0082, Fig. 2A, Suppl. Fig. 6) of equal age and sex. The antennal responses were even higher when *E. muscae* and control cadavers were in a late sporulation stage (LMM, t = 2.646, p = 0.0267, Fig. 2A). Furthermore, the complete cadaver headspace samples were attractive to male house flies in Y-tube behavioural assays at a significance level of 0.1 (exact binomial test, *p* = 0.07255, Fig. 2B, C), which supports the EAG results that male house flies are able to detect volatiles from *E. muscae* infected cadavers.

### *E. muscae* killed house fly cadavers have distinct chemical profiles of volatiles and cuticular hydrocarbons

To investigate the chemical compounds responsible for the male housefly antennal responses to conidia and sporulating cadavers, we used gas chromatography coupled to mass spectrometry (GC-MS) to analyse cuticular extracts (in hexane) of both infected and uninfected houseflies in early (3-8 hours post mortem) and late (26-28 hours post mortem) sporulation stages, equivalent to the cadavers used in the behavioural assays (Suppl. Fig. 7, 9). We observed distinct differences in the chemical profiles (Fig. 3A), which using principal component analysis (PCA) clustered according to infection status (early vs. late) and housefly sex (Fig. 3B). Distinctive compounds in cadavers sporulating with *E. muscae* compared to control fly cadavers consisted of various long-chained alcohols and esters (Fig. 3A). Many of these compounds, including canonical house fly cuticular hydrocarbons, increased from early-stage to late-stage sporulation, although the housefly host was dead throughout all sporulation stages (Fig. 3A, Suppl. Data 2).

The alkene (Z)-9-tricosene has been proposed to be the primary female house fly sex pheromone (Carlson *et al*., 1971), although other findings report that several house fly strains contain no (Z)-9-tricosene (Noorman and Otter, 2001; Darbro *et al*., 2005; Butler *et al*., 2009). We only found trace amounts of a compound with matching GC-MS properties to (Z)-9-tricosene in late sporulating males and females, and no traces in uninfected females (Fig. 3A). There was an increase in late sporulating cadavers, however, in the amounts of compounds tentatively identified as methyl-branched alkanes, most likely 2-methyloctacosane, 2- and 4-methyltriacontane and 11- and 13-methylheptatriacontane (Fig. 3A), which have previously been found to stimulate male sexual behaviour (Adams and Holt, 1987; Adams, Nelson and Fatland, 1995). The compounds 2-methyloctacosane and an unknown methyl-branched heptatriacontane have both been shown to yield small non-significant increases in mating attempts (Adams, Nelson and Fatland, 1995), while 2-, 3- and 4-methyltriacontane have previously been found to significantly increase male mating attempts (Adams, Nelson and Fatland, 1995). Altogether, these previous findings support that higher amounts of female house fly cuticular compounds increase male sexual behaviour towards female *E. muscae*-infected cadavers.

While cuticular hydrocarbons comprise the house fly sex pheromone and likely act on shorter ranges, an aerial plume of volatile compounds from the cadavers would mediate medium to long-range chemical attraction of flies to cadavers and conidia. We therefore analysed volatile compounds emitted to the surrounding air (headspace sampling) of infected and uninfected male and female fungal cadavers (Suppl. Fig. 8, 10). Infected fly cadavers had markedly different volatile profiles than uninfected control cadavers of similar age (Fig. 4A, Suppl. Data 3), while headspace from infected male and female cadavers were largely similar. Twenty-four compounds, dominated by sesquiterpenes and including an ethyl ester, ethyl octanoate, were repeatedly found in headspace from infected males and females (Fig. 4A). Interestingly, the largest single component of any *E. muscae* headspace sample, sesquiterpene 3 (Supp. Fig. 12), was also found in headspace samples of uninfected females (Suppl. Fig. 8), but not males (Suppl. Fig. 10).

**Fig. 4.**
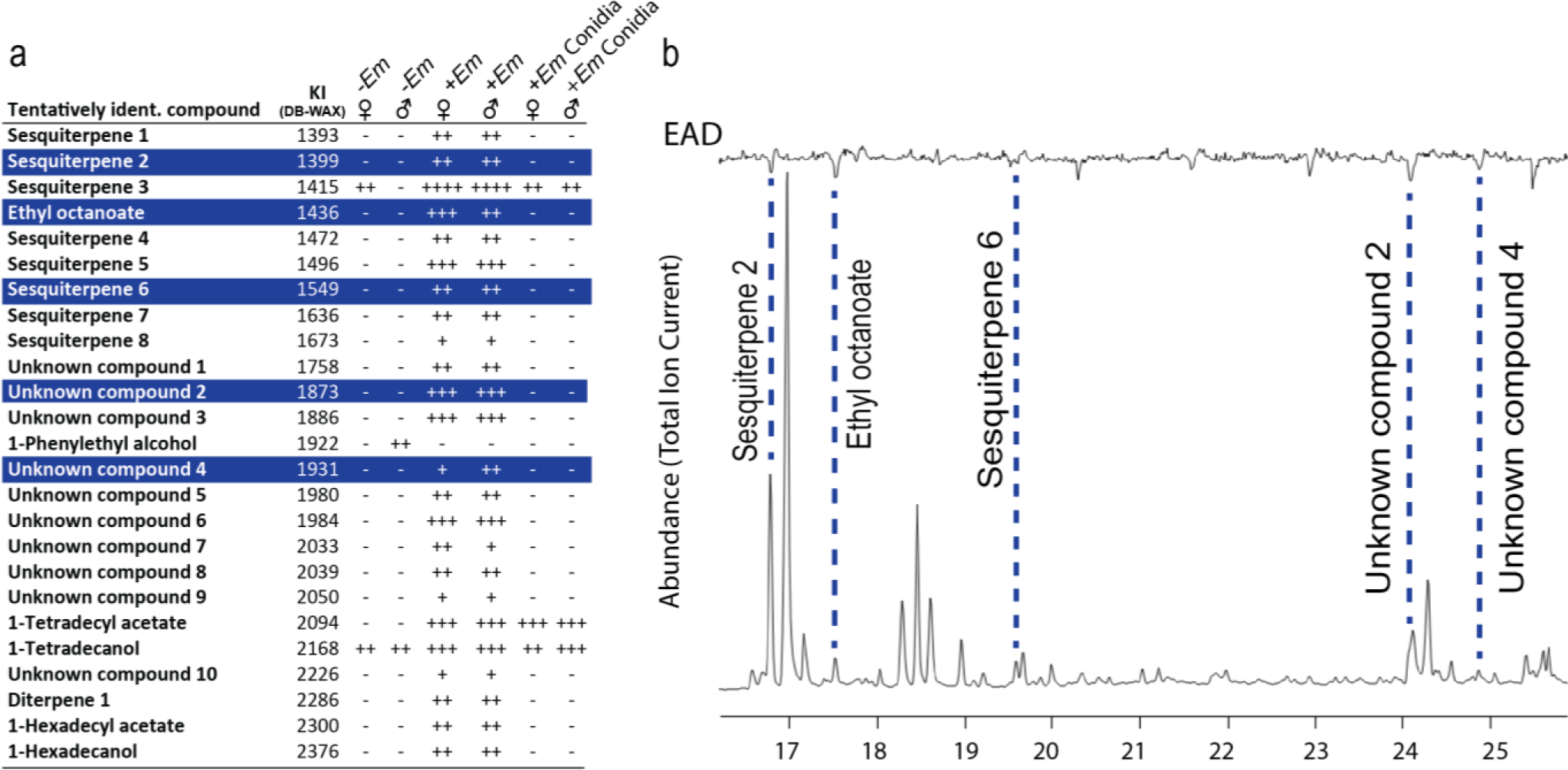
Volatile chemical profile of *E. muscae* cadavers and male antennal detection. **a**, Tentatively identified compounds found in volatile samples of uninfected (*-Em*) and infected (*+Em*) male and female cadavers sampled by collecting the ambient air (headspace) of cadavers or conidia over 24 hours. **b**, Male antennal detection of individual volatiles from infected female cadaver separated and detected using GC-EAD. Top: male antennal EAD response. Bottom: GC-FID separation of volatile compounds. Compounds marked in blue (**a**) correspond to volatile compounds highlighted with stippled lines that consistently gave an EAD response in all replicates.

Having determined that infected fly cadavers have distinct cuticular- and headspace chemical profiles, and that fly antennae elicit stronger responses to the odour of infected cadavers, we next sought to determine which compounds trigger an electrophysiological antennal response using gas chromatography-coupled electrophysiological recordings from antennae (GC-EAD, Suppl. Fig. 16). None of the long-chain hydrocarbons elicited consistent GC-EAD antennal responses, whereas ethyl octanoate (verified using synthetic standard), two putative sesquiterpenes and two compounds with unknown structure repeatedly elicited GC-EAD-responses (Fig. 4B, Suppl. Fig. 11, 13-15). Irrespective of dosage tested (20-2000 ng), ethyl octanoate did not elicit any behavioural response from male house flies when tested alone as a synthetic standard in Y-tube assays (exact binomial tests, *p* > 0.1 in all cases), indicating that ethyl octanoate alone is insufficient to trigger behavioural attraction.

### All *E. muscae* genes in the mevalonate sesquiterpene biosynthesis pathway are actively expressed in fungus-killed cadavers

Having shown that multiple volatiles from female *E. muscae* fly cadavers are detected by fly antenna and in combination are attractive to male flies, we sought to determine if the pathogenic fungus synthesize these compounds for behavioural manipulation (novel chemical attraction) or whether the male attraction is due to naturally occurring volatile chemistry in the decaying fly cadaver, for example via post-mortem house fly gene expression. Genome-wide gene expression of early (4 hours post death) and late (28 hours post death) infected and uninfected control fly cadavers, respectively, were therefore analysed for key chemical synthesis pathways (Fig. 5A). This revealed that *E. muscae* actively expressed, and in late vs early cadavers had statistically significant higher expression of several key enzymes, which are generally known to catalyse the production of precursors for characteristic bioactive esters and fatty acids (e.g., acetyl-coA carboxylase (ACC1), fatty acid synthases (FAS1, FAS2), and long-chain-fatty-acid-CoA ligase 2 (ACSL) (Eder *et al*., 2018) (Suppl. Data 4). Furthermore, in both early and late fly cadavers, *E. muscae* expressed all seven enzymes in the mevalonate pathway that synthesise the backbone isoprenoid precursors (Vranová, Coman and Gruissem, 2013) and a farnesyl-diphosphate farnesyltransferase (FDFT), required for further sesquiterpenoid and triterpenoid biosynthesis (Fig. 5B, Suppl. Data 5). Finally, we were able to identify three expressed fungal transcripts, one of which was up-regulated in 28 hour old cadavers compared to early cadavers. These transcripts showed significant homology to the yeast *Saccharomyces cerevisiae* ethyl ester biosynthesis genes *eht1* and *eeb1* (Suppl. Data 6), which are specifically involved in ethyl octanoate biosynthesis (Saerens *et al*., 2006, 2008). The expression profile of these *E. muscae* genes, in combination with a significantly lower post-mortem expression of *M. domestica* alkene biosynthesis genes in 28 hour old cadavers (Suppl. Data 7), is in concordance with *E. muscae* being solely responsible for the biosynthesis and release of the identified ethyl octanoate and sesquiterpene volatiles. Taken together this supports a novel chemical attraction of male house flies, which respond to a qualitatively improved signal from fungus-killed cadavers.

**Fig. 5.**
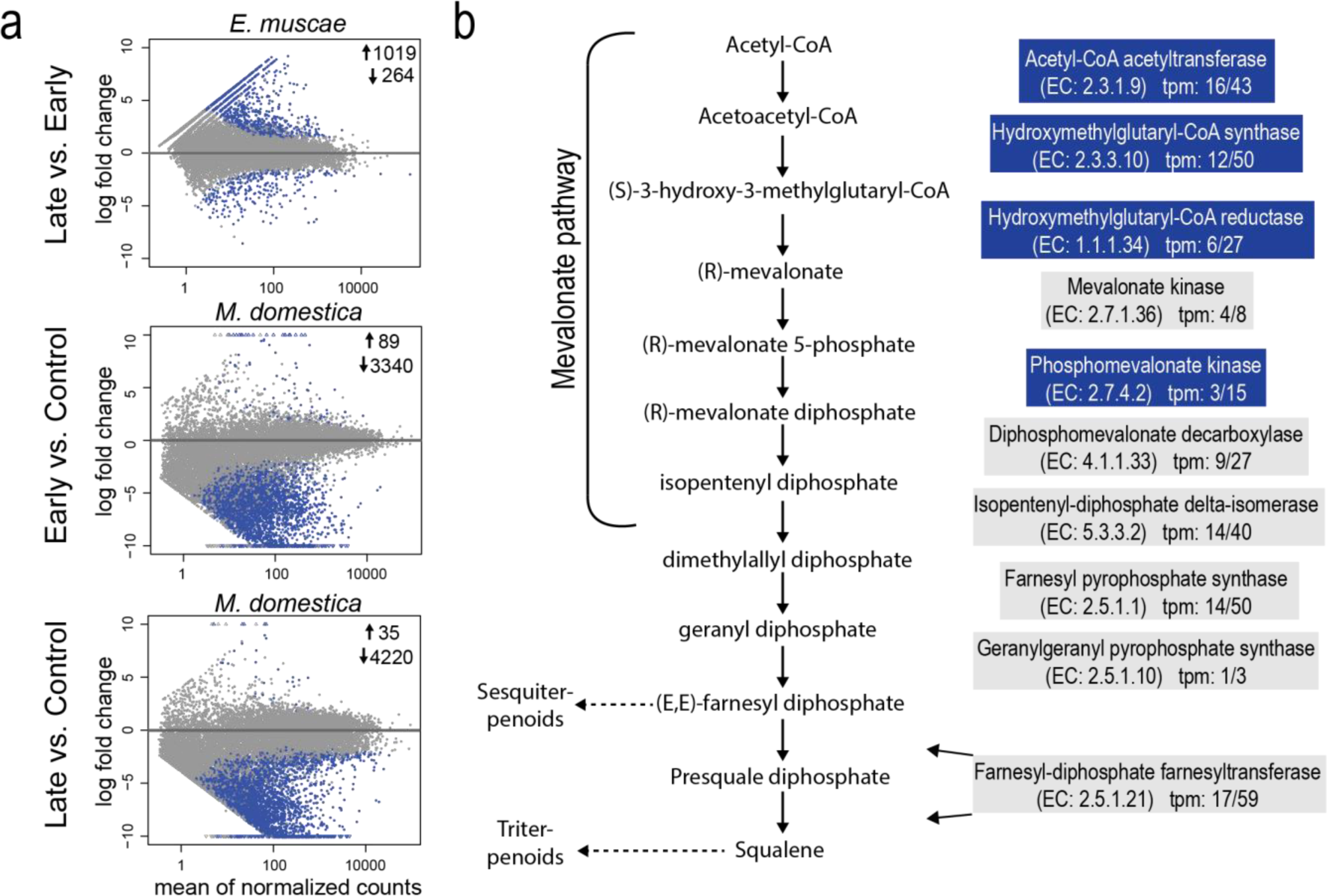
Housefly and *E. muscae* gene expression during sporulation in fungus-killed cadavers. **a**. Expressed *E. muscae* transcripts in late (26-28 hours post death) vs. early (3-8 hours post death) cadavers, and housefly expressed transcripts in early infected vs. uninfected early controls, and late cadavers vs. late controls (MA-plots). Blue dots show significantly higher or lower expressed genes (Log2 fold-change > 1; p < 0.001). **b.** All enzymes in the fungal mevalonate and sesquiterpene biosynthesis pathway were identified and expressed in *E. muscae*. Blue-coloured boxes denote enzyme-coding genes significantly higher expressed in late vs. early sporulating cadavers (Log2 fold-change > 1; p < 0.001). For each enzyme the Enzyme Commission (EC) number and expression in transcripts per million (tpm) in early / late sporulation is given.

## Discussion

Any obligate and host-specialized pathogen is in dire need of transmission to a new host. The insect-pathogenic fungus *E. muscae* represents a novel instance of volatile-mediated pathogen manipulation of host-mating behaviour (George *et al*., 2013; Trandem *et al*., 2015; Cooley, Marshall and Hill, 2018), where changes in volatile chemistry can be directly linked to changes in host sexual behaviour. Healthy males are attracted to fungus-killed cadavers and engage in courtship and mating attempts, which significantly increase infection of new host individuals and thereby ensures transmission of the fungal pathogen. The male attraction to *E. muscae* infected female cadavers cannot be explained by changes in the amount of the canonical main house fly sex pheromone (Z)-9-tricosene, or solely by visual cues from the altered size and appearance of the cadaver (Møller, 1993; Zurek *et al*., 2002). Here we show that infection with *E. muscae* induces changes in the volatile chemistry that attract house flies by both altering the levels of cuticular fly hydrocarbons and by producing several unusual volatile compounds, including several sesquiterpenes not previously associated with house flies. Sesquiterpenes have recently been found to be attractive in several other insects. For example, it has been reported that β-caryophyllene and β-elemene are attractive to *Apis cerana* (Zhang, 2018), while β-trans-bergamotene is believed to have an attractant effect on bumble bees (Haber *et al*., 2019). Terpenoids, including sesquiterpenes, are otherwise well known anti-feedants and antimicrobials (Mithöfer and Boland, 2012). For example, β-selinene is a known antifungal compound in the roots of maize, and also induced by jasmonic acid in celery (Stanjek *et al*., 1997; Ding *et al*., 2017). Generally, sesquiterpenes are not sufficiently acknowledged in the literature, most likely due to difficulties with structurally elucidating these compounds by GC-MS alone, as the mass spectra lack diagnostic peaks differentiating the many alternative structural backbones and isomers. In fact, alternative complementary approaches, such as in vivo labelling and detailed biosynthetic considerations (Könen and Wüst, 2019) or NMR-studies, requiring pure samples of >1000-fold more compound than GC-MS, are often required for the full identification of novel sesquiterpenes.

Whereas female house flies normally avoid oviposition on animal faeces colonized by harmful fungi (Lam *et al*., 2010), the volatiles found in the air surrounding *E. muscae* killed cadavers and fungal conidia were attractive to the flies. It is plausible that the volatile compounds attract male flies from a distance, but when males are within closer proximity of the cadaver they respond to the altered levels in less volatile cuticular compounds. Among the compounds that increased in fungus-killed cadavers compared to uninfected controls are specific methyl-branched alkanes similar to those known to elicit male sexual and courtship behaviours (Suppl. Fig. 17). Although the mating attempts are of shorter duration than natural matings, which normally last for more than 60 minutes (Murvosh, Fye and LaBrecque, 1964), the close physical and mechanical contact with the cadaver can trigger the active release of new infectious conidia (De Ruiter *et al*., 2019). While house flies become infected through any contact with the forcibly ejected *E. muscae* conidia (De Ruiter *et al*., 2019), the fungus-induced signalling that leads to increased male mating attempts with fungus-killed cadavers described here substantially increases the chance of infection.

We observed a significant increase in mating attempts when the cadaver was in a late sporulation stage. Close physical contact in late stage of infection increases the chance of fungal transmission because there are more infectious conidia compared to the early stage where the conidiophores are just maturing. However, when a halo of conidia was present on the surface around the cadaver in late stage of infection there was no increase in mating attempts. In spider mites infected with another entomophthoralean fungus, *Neozygites floridana*, healthy spider mites avoid conidia-covered cadavers likely because of repelling tactile cues from the conidia (Trandem *et al*., 2015). In the *E. muscae* system, however, conidia were attractive to male house flies and the negative effect on mating imply male house flies investigate or feed off the surrounding conidia rather than being stimulated to mate. The initial volatile attraction is therefore not necessarily related to sexual behaviours and could instead be linked to feeding, in particular as we observed proboscis extension towards the conidia and as they were attractive to both males and females. Alternatively, the long-range attraction could be facilitated by volatiles eliciting sexual behaviours, while the flies switch to feeding behaviour at close range or in contact with the conidia. A similar mechanism has recently been suggested for specific pollinator attraction to an Australian spider orchid, *Caladenia drummondii*, pollinated by solitary thynnine wasps (Phillips, Bohman and Peakall, 2021). Behavioural attraction of males towards volatiles of late cadavers were highly depend on the time of day, which is suggestive of a relation with mating behaviours, which is known from other fly species such as *Drosophila suzukii* to be highly influenced by time of day (Revadi *et al*., 2015).

Feeding and sexual behaviour is tightly linked in *D. melanogaster* (Spieth, 1974; Grosjean *et al*., 2011), which makes it difficult to distinguish whether the attracted house fly males are lured in from a distance due to mating cues or feeding cues (i.e. sexual or food mimicry, respectively).

*E. muscae* can induce epizootics in house fly populations (Mullens, Rodrigues and Meyer, 1987), and previous findings indicate that horizontal transmission can occur via conidial exchange between healthy males exposed to conidia and healthy females (Watson and Petersen, 1993). An increase in male sexual behaviour towards sporulating cadavers may thus serve the dual purpose both to infect the focal male fly, but also as a viable option for wider fungal transmission throughout a population. House flies have an emerging role in animal feed as house fly larvae (van Huis *et al*., 2020), but especially under unsanitary conditions house flies also act as mechanical vectors for more than 100 disease-causing human pathogens (Khamesipour *et al*., 2018). This ambiguous role leads to situations where regulation of house fly populations may be necessary. The findings presented here may thus have potential for the discovery of novel semiochemical house fly-specific attractants or pheromones that could be used in pest control.

While the EAD-active sesquiterpenes emitted by *E. muscae* cadavers could not be identified in detail, ethyl octanoate was verified using a synthetic standard. This semiochemical is attractive to *D. melanogaster* (Schiabor, Quan and Eisen, 2014) and the tephritid fruit fly *Anastrepha ludens* (Robacker, Warfield and Flath, 1992; Malo *et al*., 2005) and is generally associated with fruits (Riu-Aumatell *et al*., 2004) and yeast fermentation (Gómez-Míguez *et al*., 2007; Eder *et al*., 2018). The volatile organic compounds produced by *E. muscae* clearly serve a manipulative behavioural function to attract healthy susceptible hosts, but have likely evolved from compounds produced for other purposes (Biedermann, De Fine Licht and Rohlfs, 2019). Such precursors could be structural compounds, compounds with an integral function for the fungus’ survival, or compounds that protect the cadaver from biotic and environmental factors. The functional adoption of ancestral compounds as fungus-emitted volatiles thus represent the evolution of an extended phenotypic trait (Dawkins, 1981) that exploit male flies’ willingness to mate and benefit the fungus by altering the behavioural phenotype of uninfected healthy male host flies. As such, this is one of the first descriptions of a behaviour-manipulating pathogen that extends beyond the manipulation and death of the focally infected host, to also manipulate the behaviour of healthy individuals.

## Methods

### Insect culture

House flies (wildtype *Musca domestica*, strain 772a) were obtained as pupae from the department of Agroecology, Aarhus University, Denmark. After eclosion, flies were sexed and separated into sexes within 24 hours in cages of around 30-50 flies. Adult house flies were kept in cylindrical plastic containers (diameter: 7.5 cm, height: 8 cm) with a net covering the top. Here, they were continuously supplied with a diet consisting of 1:1 (V/V) skim milk powder and sugar, and water supplied from a 15-mL centrifuge tube inserted into the side of the container sealed with a cotton ball wick. They were kept at a 16:8 hour light/dark rhythm at an average temperature of 21 ±1 ℃.

### Fungal cultures

House flies were infected with *Entomophthora muscae* (isolate no.: KVL21-01, University of Copenhagen, Section for Organismal Biology Entompathogenic fungus culture collection) originally isolated from a house fly caught in a cow stable (“Birkedal”, 55.835571, 12.154350) and the fungus were continuously maintained inside house fly hosts as previously described (Hansen and De Fine Licht, 2017). Once exposed to a *E. muscae* conidia shower, infected flies were moved to an individual incubator with a constant temperature of 19,5 ℃. Here, a reverse light-dark rhythm consisting of 14:10 hours light/dark was applied, with the light ending at 10:00 am each morning. Six to seven days post *E. muscae* exposure, infected flies were checked and cadavers exhibiting clear *E. muscae* infection were collected at the end of the photoperiod. As *E. muscae* induces death synchronized with the end of the photoperiod in infected flies (Krasnoff *et al*., 1995), the end of the photoperiod also corresponds to time of death and the majority of these cadavers were only on the verge of sporulation when they were collected (seen by visible conidiophores extruding from the cadaver).

### Mating activity experiment

In order to determine male house fly sexual attraction towards *E. muscae* infected cadavers, mating activity experiments were performed. Here, an uninfected virgin male was anaesthetized by cooling at 5 ℃ for 2 minutes and placed within a clean, EtOH-rinsed glass Petri dish arena (diameter: 90 mm, height: 16 mm), together with an infected or an uninfected control cadaver, fixated to the bottom with petroleum jelly. The experiments were conducted at a room temperature of 21 ± 1 ℃. Cadavers were divided into two groups: early killed (equivalent to an early sporulation stage), and late killed (equivalent to a late sporulation stage). Early killed cadavers were used within 3 to 8 hours post mortem, equivalent to most of the scotophase. Late killed cadavers were placed in a chamber with 85% RH humidity immediately after death to prevent desiccation, and used in experiments between ∼ 25 to 30 hours post mortem. All uninfected house flies were killed by freezing for 6 minutes at -24 ℃ and *E. muscae* infected females and males were killed by *E. muscae* infection and showed characteristic white, fungal bands of conidiophores. All mating experiments were performed in the same room as rearing. For mating experiments, male-female and male-male trials were conducted from 10:48 am to 17:53 pm and from 09:24 am to 17:49 pm respectively.

The glass Petri dish arena was placed on a printed sheet of paper which denoted the exact middle of the arena, as well as an inner circle with a 16 mm radius around the cadaver. A video camera was placed 12 cm above the glass Petri dish arena. The behaviour of the male fly was filmed and monitored for 40 minutes and behaviors related to sexual attraction were noted in the software BORIS v. 6.2.4 (Friard and Gamba, 2016). Here, the number of mating attempts, time spent attempting mating, number of physical contacts, and time spent in the vicinity of the cadaver, were noted. The standard definition of house fly mating behaviour, a “mating strike”, was used as previously described (Murvosh, Fye and LaBrecque, 1964). A physical contact was defined as any physical contact between the male and the cadaver that was not a mating attempt nor was directly associated with one. Such physical contact varied from touching of legs or wings to if the subject crawled entirely over the cadaver. A new physical contact would be noted after complete dissociation of male and cadaver followed by a new touch. Time spent in the vicinity was measured as the time any body part of the male spent inside the 16 mm radius of the cadaver.

In order to determine successful *E. muscae* infections from associating with cadavers during assays, the living male subjects were anaesthetized with CO_2_ and incubated individually in clean fly cages on a 1:1 skim milk powder sugar diet, supplied with clean water, immediately after each 40 minutes of observation. Flies were monitored daily until death and checked for visible fungal growth in the intersegmental membrane abdomen, characteristic of *E. muscae* infection.

### Mate choice experiment

Mate choice experiments were performed as a two-choice assay in polystyrene Petridishes (diameter: 92 mm, height: 16 mm). A cadaver of each tested treatment was fixated with its ventral side of the thorax to the bottom in each side of the arena with petroleum jelly. Cadavers were placed so the heads faced the side and the abdomen faced the center of the arena. As in the mating activity experiments, a video camera was placed 12 cm above the arena. A healthy virgin male was anaesthetized by cooling at 5 ℃ for 2 minutes and placed in the center of the arena, where after the male subjects were filmed for 40 minutes. The number of mating attempts (as previously defined) towards either cadaver were noted in the software BORIS v. 6.2.4 (Friard and Gamba, 2016).

### Conidia attraction experiment

To determine attraction of *E. muscae* conidia, we performed a two-choice conidia attraction cage experiment. Here, two sticky trap discs had been exposed to a conidia shower from three cadavers of one sex, that were either *E. muscae* infected or uninfected freeze-killed controls (-24 ℃, 6 minutes) for 24 hours in 85% RH humidity. The sticky trap discs had been placed in the bottom of a polystyrene Petri dish (diameter: 92 mm, height: 16 mm), with the three cadavers fixed to the lid. After 24 hours, the lid and cadavers were removed and replaced with a new lid with a 1 cm diameter hole in the center. Both treatments of Petri dish traps (conidia and control) were wrapped in Parafilm, not covering the hole to eliminate visual cues, and placed inside a plexiglass cage (dimensions: 30 x 30 x 30 cm). Four virgin flies of the same sex (5-20 days old) were released in the cage and allowed to choose between traps for 24 hours. After 24 hours, the number of flies in each trap were counted and a preference index calculated (Quan and Eisen, 2018):

A = total number of flies in trap with conidia, B = total number of flies in trap with control:

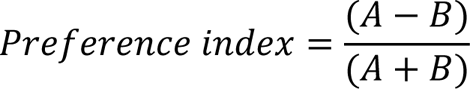

A positive PI indicates a preference for *E. muscae* conidia, whereas a negative PI indicates preference for the control. Flies that did not make a choice to either trap were excluded, as were trials where none of the four flies made a choice.

### Gas Chromatography-Mass Spectrometry (GC-MS) of cuticular compounds

House fly samples were prepared in the exact same way as for the mating activity experiment and encompassed males and females in early (3 – 8 hours post mortem) and in late sporulation stages (25-30 hours post mortem) and corresponding controls (n = 5 of each treatment, all flies 8-10 days old). Conidia-samples were obtained by inserting individual sporulating house fly female (n = 2, 13 days old) and male (n = 5, 9 days old) cadavers into cut pipette tips, so that the abdomen protruded from the end. This pipette tip was then gently inserted into a 2-mL glass vial and placed in a humid chamber with 85% RH for 24 hours. This allowed the conidia to be discharged from the abdominal conidiophores into the vial without contact to the fly cadaver. House fly cuticular compounds and hexane-soluble fungal metabolites from individual flies, cadavers or conidia were extracted in 2-mL vials with 500 µL *n*-hexane (for liquid chromatography LiChrosolv®, Sigma-Aldrich) for 5 minutes. Thereafter the hexane extract was transferred to a new vial and concentrated under a steady stream of N_2_ to a total volume of 40 µL before analysis.

Extracts were injected into an Agilent 6890N GC equipped with a DB1 MS-UI column (60 meter length, 0.25 mm inner diameter, 0.25 µm film thickness, Agilent Technologies) coupled to a 5975 inert Mass Selective Detector (Agilent Technologies). Splitless injection (0.5 min) was applied (injector temperature 325 ℃). The oven was programmed from an initial temperature of 200 ℃ for 2 minutes and a ramp up of 8 ℃/min to 340 ℃ and held for 15 minutes. Helium was used as the carrier gas with a linear velocity of 35 cm/s. The electron ionisation mass spectra were recorded at 70 eV and samples were analysed in MSD ChemStation v. D.03.00.611 (Agilent Technologies). Retention times were related to an injected Kovats linear alkane mixture of carbon chain length C8-C40 and linear retention indices for each peak were calculated (van Den Dool and Dec. Kratz, 1963). For tentative compound identifications, we used library searches against a NIST14 library and by comparing Kovats indices, mass spectra and diagnostic ions in previously published analyses (Nelson, Dillwith and Blomquist, 1981; Bagnères and Morgan, 1990; Stránský *et al*., 1992, 2006; Carlson, Bernier and Sutton, 1998; Mpuru *et al*., 2001; Gulias Gomes, Trigo and Eiras, 2008; Zhang *et al*., 2010).

### Gas Chromatography-Mass Spectrometry (GC-MS) of volatile compounds

Headspace samples were collected in a dynamic headspace sampling (aeration) of 20-22 hours. The aeration was collected from five cadavers of each sex, either actively sporulating or freeze-killed control cadavers, and blank samples without cadavers. The cadavers were placed in a glass Petri dish and enclosed in a cooking bag (Toppits). The outlet of an electrical air pump (KNF Neuberger, model PM 10879-NMP) was connected with silicone tubing, pushing air through a charcoal filter, tightly connected with the bag through a small opening. Similarly, at the opposite side of the bag, a filter made of Teflon-tubing and Porapak Q 50/80 adsorbent (Markes International) was tightly connected and joined with the pump air inlet to a closed-loop headspace collection system. Airflow was adjusted to 0,75 L/min for all samples. Adsorbed headspace samples were subsequently eluted from the Porapak filter with *n*-hexane (2 x 500 µl) into 2-mL glass vials and concentrated under a gentle stream of N_2_ to a total volume of 40 µL before analysis. Samples (2 µL) were injected into a 7890B GC/5977A GC/MSD system (Agilent Technologies) with a DB-Wax capillary column (60 m length, 0.25 mm inner diameter, 0.25 µm film thickness). Splitless injection (0.5 min) was applied (injector temperature 225 ℃). The oven was programmed from an initial temperature of 30 ℃ for 3 minutes and a ramp up of 8 ℃/min to 225 ℃ and held for 10 minutes. Helium was used as carrier gas with a linear velocity of 35 cm/s. The electron ionisation mass spectra were recorded at 70 eV, and compounds were tentatively identified based on library searches against the NIST14 library and Kovats indices from an injected alkane mixture of carbon chain length C8-C40 (van Den Dool and Dec. Kratz, 1963). Ethyl octanoate was identified by injection of a synthetic standard. Compounds found in at least three out of five samples for each treatment and not found in blank control samples, were included.

### Electrophysiological recordings (EAG & GC-EAD)

Electroantennography (EAG) and Electroantennographic Detection (GC-EAD) recordings were performed to measure male house fly antennal responses to infected and uninfected cadavers, and to identify key fungal odour components that elicited antennal responses. Electrophysiological recordings were performed on antennae of whole male house flies mounted in a cut pipette tip. Pulled glass capillaries with a silver wire and filled with Ringer solution (Merck Millipore, product no.: 115525) were placed on the tip of the funiculus and in the eye as recording and ground electrodes, respectively (Suppl. Fig. 16). Antennal responses were digitized with an IDAC-2 system (Syntech, Kirchzarten, Germany) and measured in software GcEad 2014 v 1.2.5 (Syntech). Carbon filter-purified air was humidified and delivered at 1,5 L/min to the antennae via a glass tube.

For EAG recordings, pulse stimuli (0.5 second puffs) were given with glass Pasteur pipettes containing the odorant on a filter paper (100 ng ethyl octanoate, n = 10), an entire fly, or *E. muscae* conidia. Fly puff measurements were performed with fly cadavers with treatments similar to the mating activity experiment early sporulation (n = 6), late sporulation (n = 10), or with a living, uninfected fly (n = 10). Pasteur pipettes for delivering puffs of conidia headspace or its respective control were prepared by placing an infected or uninfected, freeze-killed fly inside the Pasteur pipette for 24 hours and allowing it to sporulate at 85% RH. The cadaver was then removed, leaving only conidia inside (n = 10). For puff stimulation, the tips of the glass pipettes were gently inserted into an opening of the side of the glass tube supplying the airstream of the EAG system onto the fly antennae.

For GC-EAD recordings, cuticular extracts (2 µL) or headspace samples were injected manually into a 7890A GC System (Agilent Technologies) with a DB-Wax capillary column (30 m length, 0.25 mm inner diameter, 0.25 µm film thickness) as the stationary phase. Splitless injection (0.5 min) was applied (injector temperature 215 ℃). The oven was programmed from an initial temperature of 30 ℃ for 3 minutes and a ramp up of 8 ℃/min to 225 ℃ and held for 20 minutes. Helium was used as carrier gas with a linear velocity of 45 cm/s. The GC was coupled to an EAD and a flame ionization detector (FID). Retention indices of any response-eliciting peak were determined based on injected linear alkane mixture of carbon chain length C8-C20 and compared to that of the GC-MS headspace injections. Ethyl octanoate was injected as a synthetic standard to confirm antennal EAD-activity in cases where there was no response. For both EAG and GC-EAD, responses were determined by measuring the amplitude of depolarization in millivolt (mV) elicited on the antenna (n = 3 biological replicates with 2-3 technical replicates on each housefly) by the individual treatment different to baseline.

### Y-tube olfactometer choice test

Male house fly attraction to infected or uninfected headspace samples were tested in a glass Y-tube olfactometer. The olfactometer had a 17 cm long base and 9 cm long arms at 90° angle with an inner diameter of 2 cm. Each side was connected to a glass cylinder (5 cm long, 1.5 cm internal diameter), which served as an odour-release compartment and was separated from the air supply with a piece of metal netting. A steady stream of humidified, charcoal-filtered air was supplied to each arm at 0.4 L/min with an air pump (KNF Neuberger, model NMP 830 KNDC). The glass Y-tube was washed, rinsed with ethanol (70%) and baked at 120 ℃ for 2 hours before each round of experiments. Headspace samples were collected with an active headspace sampling method (aeration) and eluted with *n*-hexane similar to GC-MS analysis of headspace. 10 µL infected or uninfected headspace samples (10 µL) were loaded onto a filter paper (1.5 cm x 1 cm), and used in the Y-tube immediately after the solvent had evaporated. Odor sources were only used for a single choice test and the positions of each odor was switched after each test. Healthy, unmated flies were allowed to walk from their cage individually into *Drosophila* polypropylene tubes (WVR) with a cotton plug and allowed to acclimate for 30 minutes, before they were introduced at the base of the olfactometer. A response choice to uninfected control- or infected house fly headspace was made once the male entered past 5 cm in one arm within 3 minutes. All Y-tube choice tests were made at room temperature under diffuse dim light. We observed a strong effect of time of day, where only experiments conducted in the afternoon (2 PM – 6 PM) yielded a behavioural response, and data for this time-period were therefore analysed by calculating a preference index per group of flies that made a choice per hour for each day the test was conducted.

### RNA Sequencing and transcriptomic analysis

Five early sporulation stage cadavers, five late sporulation stage cadavers, five early uninfected control cadavers, and four late uninfected control cadavers, all unmated female flies that had been prepared similarly to cadavers in the mating activity experiment, were collected. All cadaver samples were 10-11 days old at the time of death. The cadavers were snap-frozen in liquid N_2_ and stored at -80 ℃ until extraction. For extraction, samples were crushed with a mortar and pestle and further homogenized with glass beads in a TissueLyzer (Qiagen). Total RNA was extracted using phenol/chlorophorm/isoamylalcohol phase-separation followed by a GeneJET RNA Purification Kit (Thermo Scientific™), according to the manufacturer’s instructions. PolyA-mRNA library preparation and 100 bp paired-end un-stranded DNBseq sequencing were conducted by BGI Europe A/S. For quality control and filtering, reads with low quality were filtered using a maxEE value of 2, while the first 10 bp of each read were trimmed, and all reads had a minimum length of 35 bp using function “filterAndTrim” in the R-package “dada2” (R version 4.0.3). FastQ files were visually inspected before and after trimming with FastQC (Babraham Bioinformatics, 2019). Reads were mapped to a concatenated reference consisting of house fly (*Musca domestica,* ncbi version 2.02, RefSeq: GCF_000371365.1) and *E. muscae* transcripts (European Nucleotide Archive (ENA) at EMBL-EBI, accession no. ERZ2299657) using the software kallisto (Bray *et al*., 2016) version 0.46.1. Differential expression analysis was conducted in R version 4.0.3 using the package “DEseq2” requiring a fold-change > 1 and a p-value < 0.001 adjusted for multiple testing to designate significant differential expression per gene. Samples were analysed pairwise as infected early vs control early and infected late vs control late for house fly genes, whereas *E. muscae* transcripts were compared between infected late vs infected early.

### Statistical analysis

All statistical analysis of data were performed in R version 4.0.3. The mating activity experiment and choice mating experiment were analysed similarly, with the house fly mating attempts using treatment as a descriptor with a Kruskal-Wallis rank sum test. Pairwise comparisons were performed with a Dunn’s test for multiple comparisons with a Bonferroni adjustment.

In the conidia attraction experiments, only flies caught in a trap were used for analysis. In order to focus on the attraction towards conidia alone, the flies that had not “chosen” a trap were excluded. Number of counted flies were analysed with a logistic regression. Choices towards either trap were described as a function of trap treatment (conidia or control), the choosing sex, the sex of the conidia-discharging cadaver and house fly age as a random factor. Y-tube experiments were grouped based on the time of the day the experiment was performed and analysed using the exact binomial test.

Signal amplitudes of electroantennographic recordings were log-transformed to fit normality and tested with a Shapiro-Wilks test to meet the assumptions for parametric tests. The amplitudes obtained from stimulation with cadavers or conidia were analysed with a linear mixed model and milli-Volts (mV) were described as a function of treatment and with the mounted fly as a random factor. Tukey’s Posthoc tests were performed for pairwise comparisons with a Bonferroni correction. Statistical analyses were performed using R packages lme4 and emmeans.

Principal component analysis was calculated on fourth-root transformed total ion chromatogram (TIC) abundance counts of compounds found in cuticular hexane extracts and extracts from conidia, using packages “FactoMineR” and “Factoextra” in R, version 4.0.3. Compound peaks that in infected fly samples contained co-eluting fungus-derived compounds were excluded from analysis from all sample types.

## Data availability

The RNAseq data that support the findings of this study are available from the National Center for Biotechnology Information’s Short Read Archive (SRA) (BioProject ID: PRJNA758214). All other data are provided in the supplementary material linked to this article.

## Code availability

No custom code was generated for this study.

## Acknowledgements

A.N. and H.H.D.F.L. were supported by a grant (no. DFF - 7014-00188) to HHDFL from the Independent Research Fund Denmark.

## Author contributions

A.N. and H.H.D.F.L. conceived of the study. A.N., H.H.D.F.L., P.G.B., and A.B.J. designed the study. A.N. performed *E. muscae* in-vivo rearing, behavioural assays and analysis, chemical sample collection, RNA extraction. A.N, P.G.B., and C.A.K. analysed chemical data. A.N., and H.H.D.F.L. analysed RNAseq data. A.N. and H.H.D.F.L. wrote the initial draft of the paper. A.N., H.H.D.F.L., P.G.B., A.B.J., B.B., C.A.K. contributed to interpreting the data and editing subsequent drafts of the manuscript.

## Supplementary material

**Supplementary figure 1.**
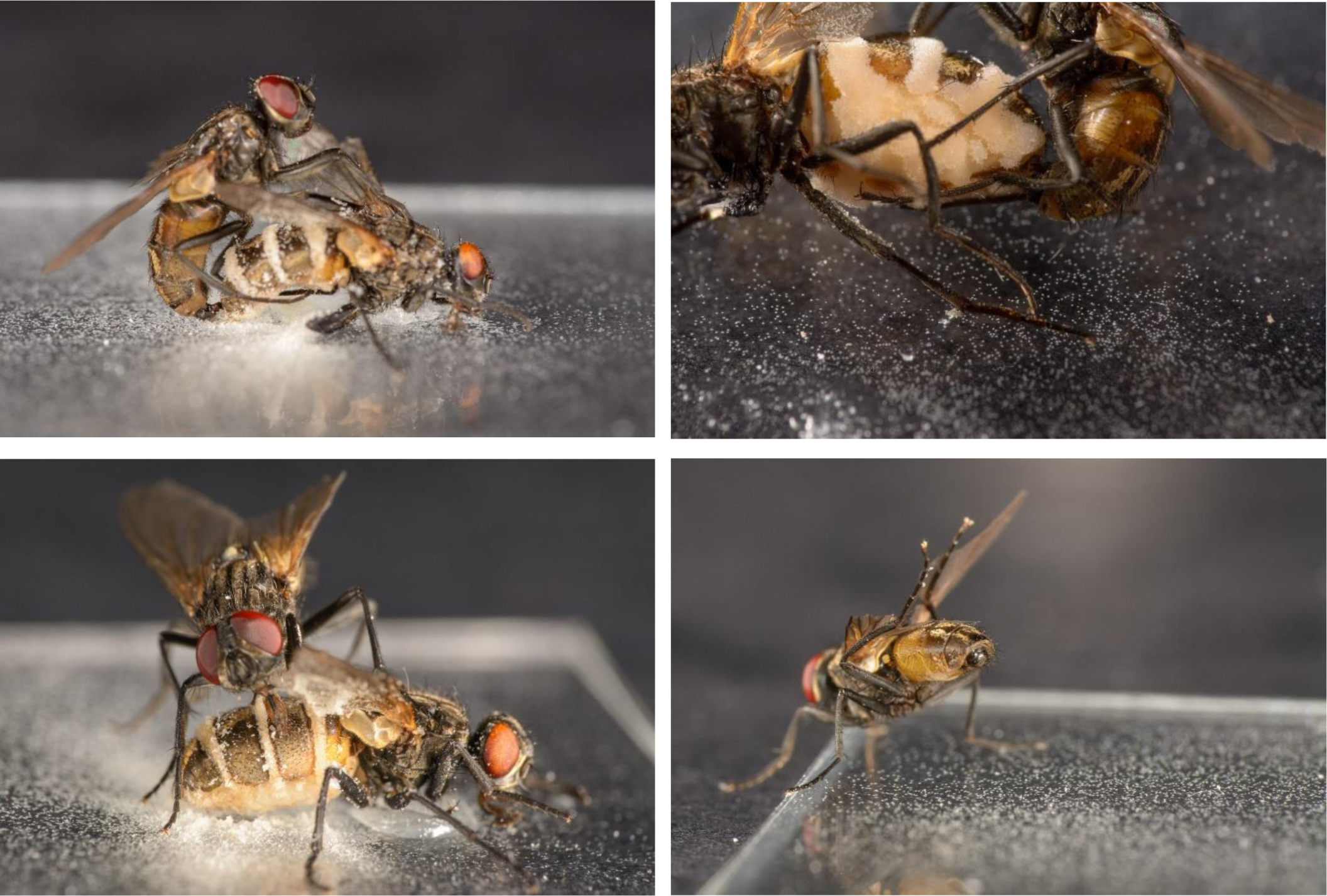
Housefly behavior towards *E. muscae* sporulating cadavers. **a**. A healthy, male housefly attempting copulation with an *E. muscae* sporulating housefly. **b.** Male attempting copulation with a sporulating female housefly. The male visibly extends his aedeagus to engage in mating. **c.** A housefly extending its proboscis and feeds on the sporulating cadaver. This behavior was exhibited towards both cadavers and conidia alone. The cadaver is fixated to the glass surface with a small amount of petroleum jelly (Vaseline) on its thorax. **d.** Healthy male housefly engaging in grooming behavior after being in contact with cadaver and conidia. Here, several conidia has already attached themselves to legs and abdomen. In all photos conidia can be seen as white powder primarily on surfaces and the cadaver, but also on the male’s abdomen and legs. Photos: Fillipo Castelucci.

**Supplementary figure 2.**
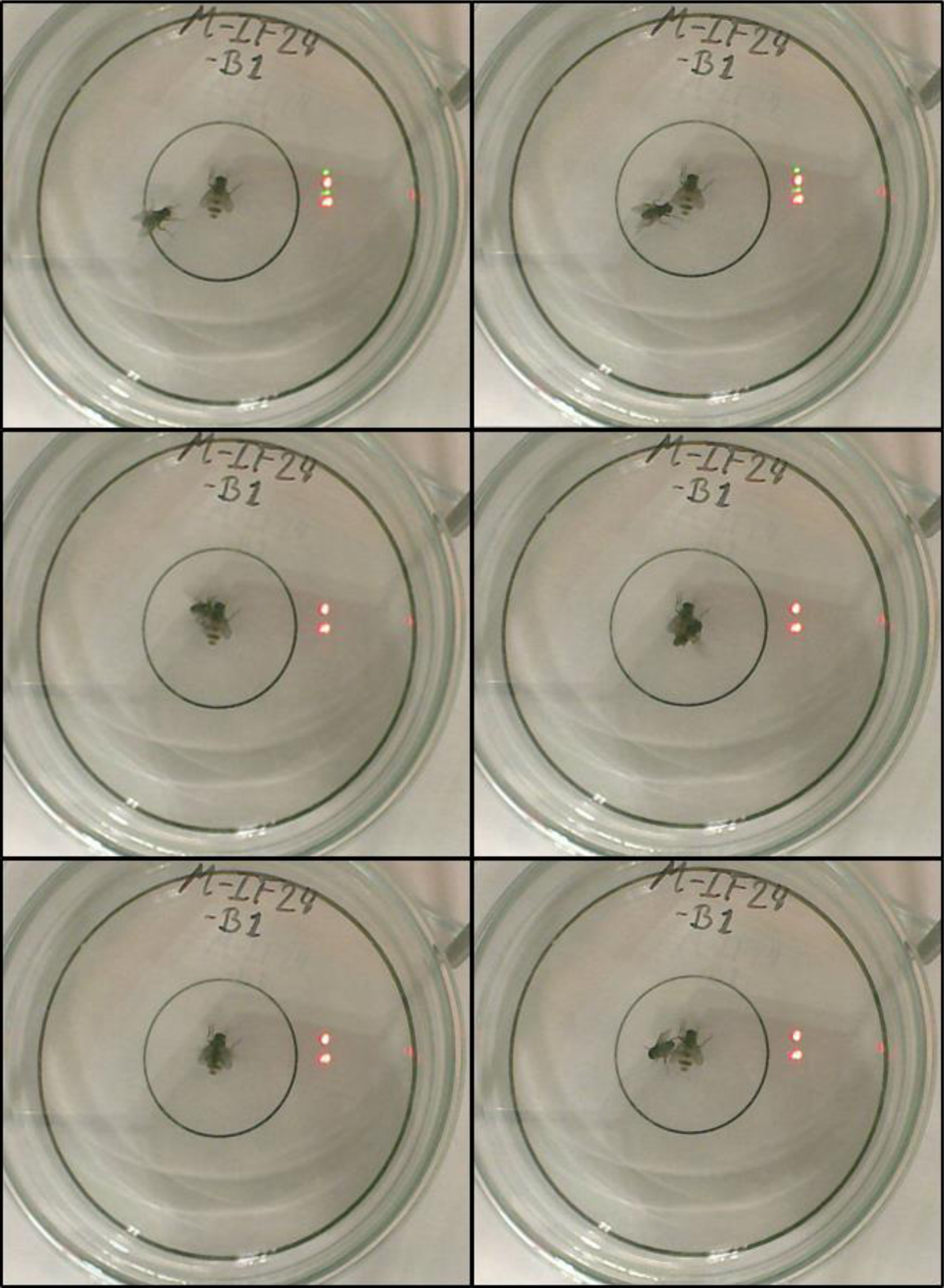
A mating attempt as noted in the mating activity experiment. A single cadaver is fixated in the center of a glass arena (Petri dish). The arena is placed on top of a printed sheet of paper, denoting the center (cadaver), an inner ring (radius = 16 mm), which denotes when the subject is “in vicinity” and an outer ring to indicate the outer borders of the arena. After acclimating to the arena, the male will usually first interact with the cadaver by physical contact. A mating attempt occurs when the male approaches the cadaver, usually from the sides or rear, and jumps onto the cadaver, accompanied by excessive wing flickering. Then he attempts to connect his aedaegus with the tip of the cadaver’s abdomen.

**Supplementary figure 3.**
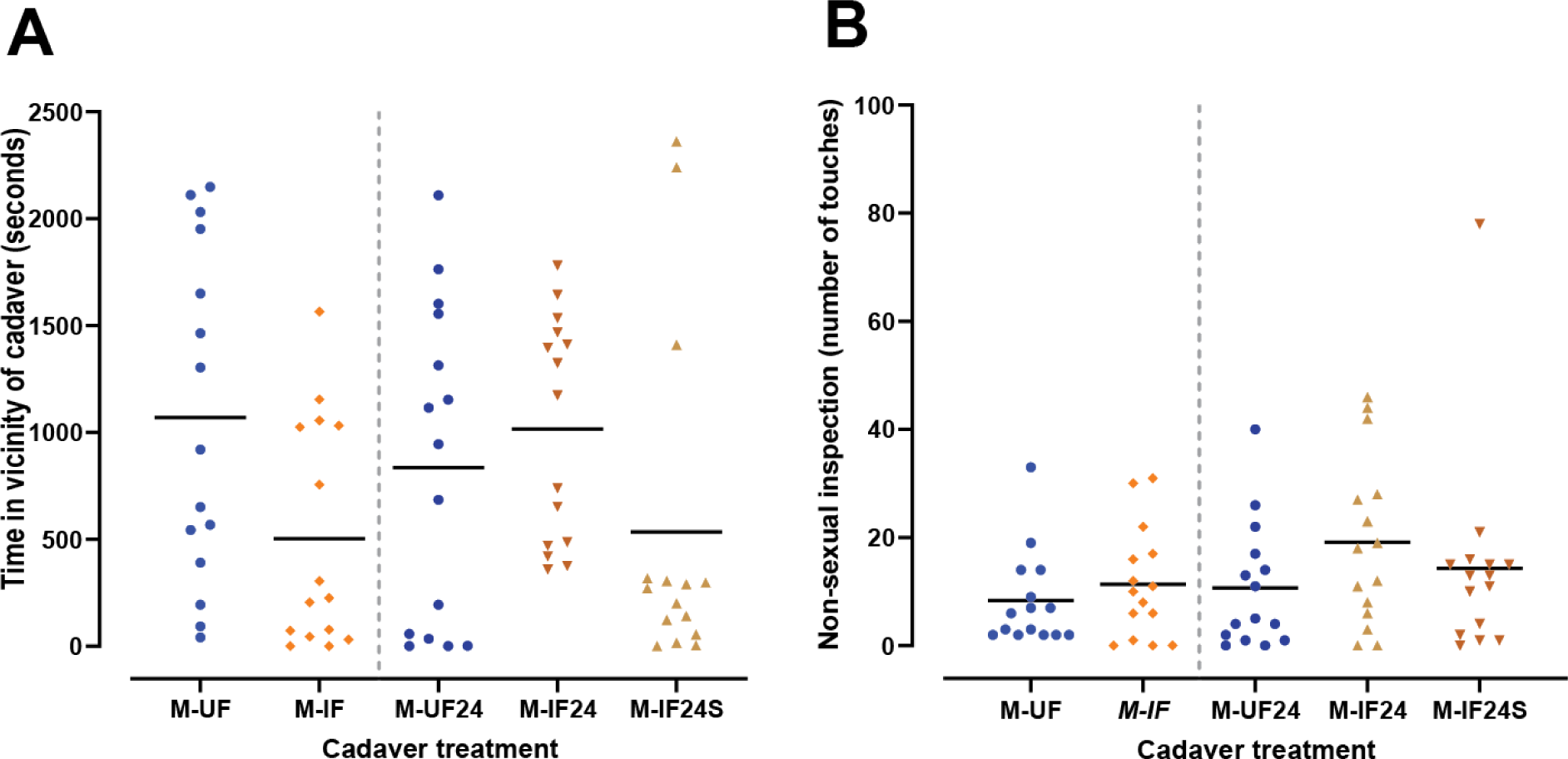
Male behaviour towards female cadavers. **A.** Time spend in vicinity of cadaver (radius: 16 mm). Male on uninfected female early control (M-UF, n =15), M-IF: Male on infected female, early sporulation stage (M-IF, n=15), Male on uninfected female, late control (M-UF24, n =15), Male on infected female, late sporulation stage (M-IF24, n = 15), Male on infected female, late sporulation stage with spores around the cadaver (M-IF24S, n = 15) **B.** Number of non-sexual physical cadaver inspections.

**Supplementary figure 4.**
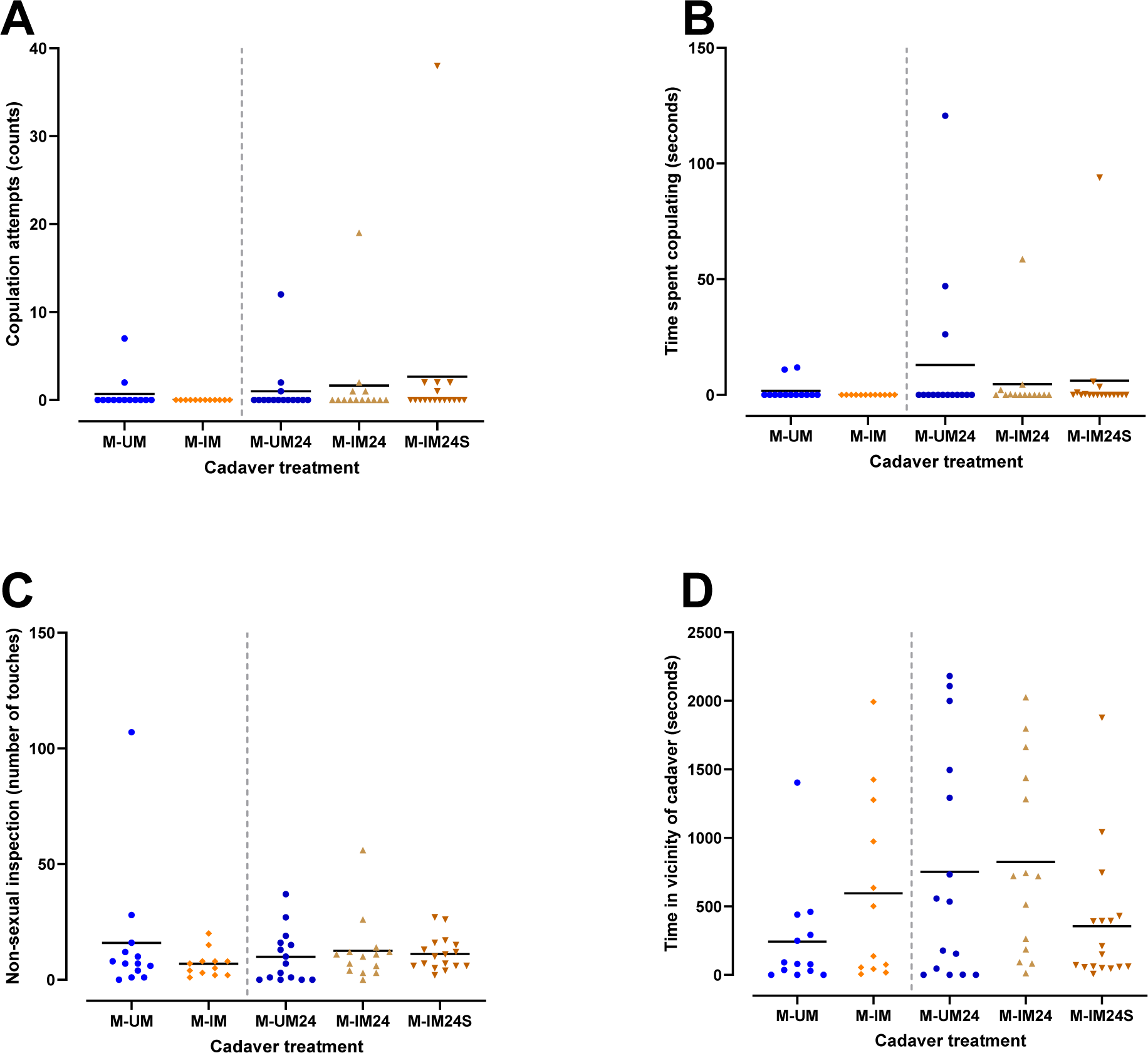
Male on male mating behavior. **A.** Number of mating attempts measured per trial. Male on uninfected male early control (M-UM, n =13), M-IM: Male on infected male, early sporulation stage (M-IM, n=12), Male on uninfected male, late control (M-UM24, n=15), Male on infected male, late sporulation stage (M-IF24, n=14), and Male on infected male, late sporulation stage with halo of conida around the cadaver (M-IF24S, n=17) **B.** time spent mating (seconds) **c.** Non-sexual inspections (counts of touches), **d.** time spent in vicinity (radius: 16 mm) of the cadaver.

**Supplementary figure 5.**
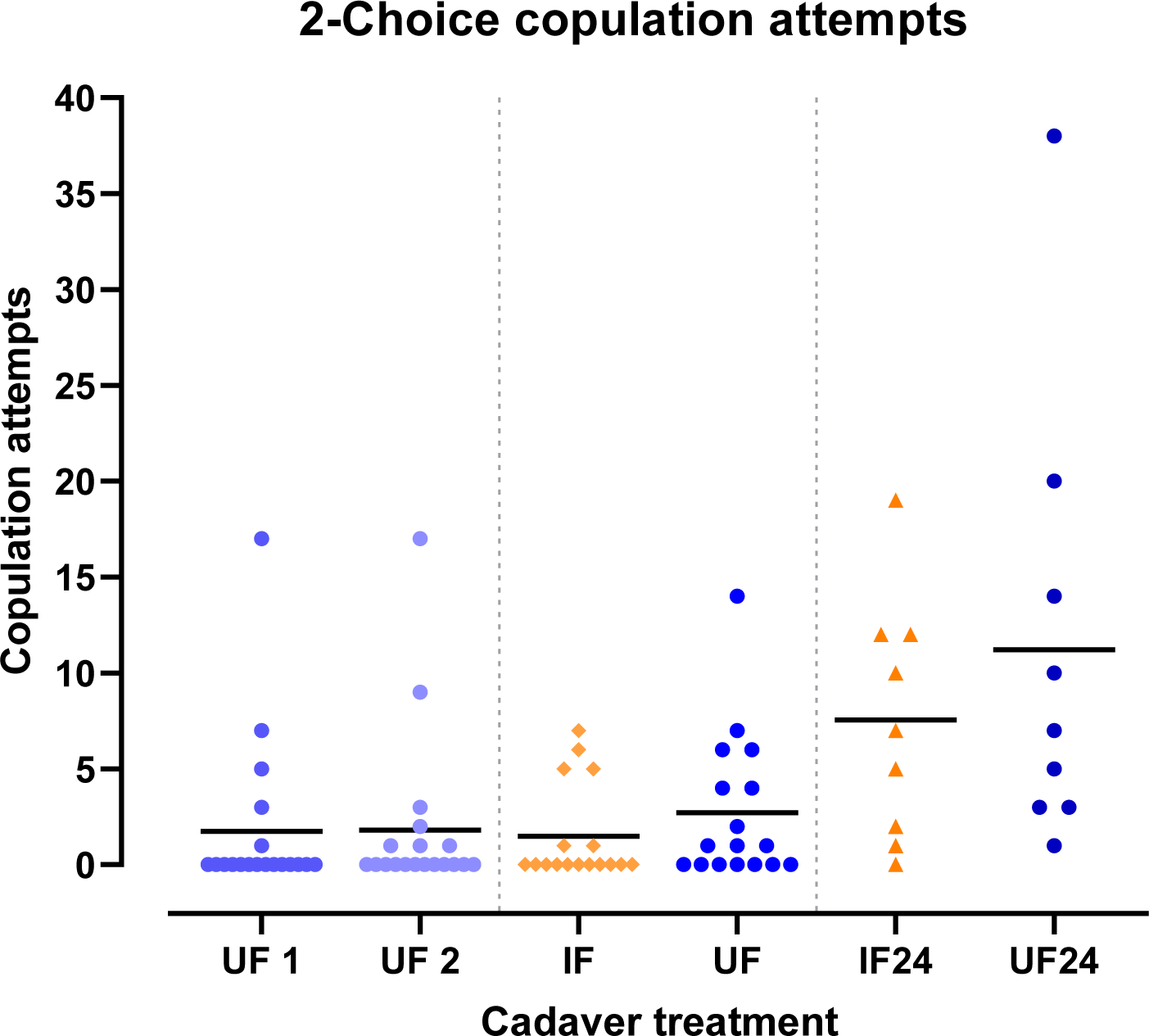
Petri dish 2-choice experiment. One male were allowed to choose between two female cadavers. In the first subset (left), two uninfected, freeze-killed controls were placed and named UF 1 & UF 2 (n = 19). In the second subset (middle), a male was allowed to choose between an uninfected female control (UF), and an *E. muscae* cadaver (IF) in an early sporulating stage (n = 17). In the last subset (right), a male was allowed to choose between a freeze-killed, uninfected control (UF24) and an *E. muscae* sporulating cadaver (IF24) in a late sporulation stage (n = 9). There was no significant differences in the amount of copulation attempts in either experiment.

**Supplementary figure 6.**
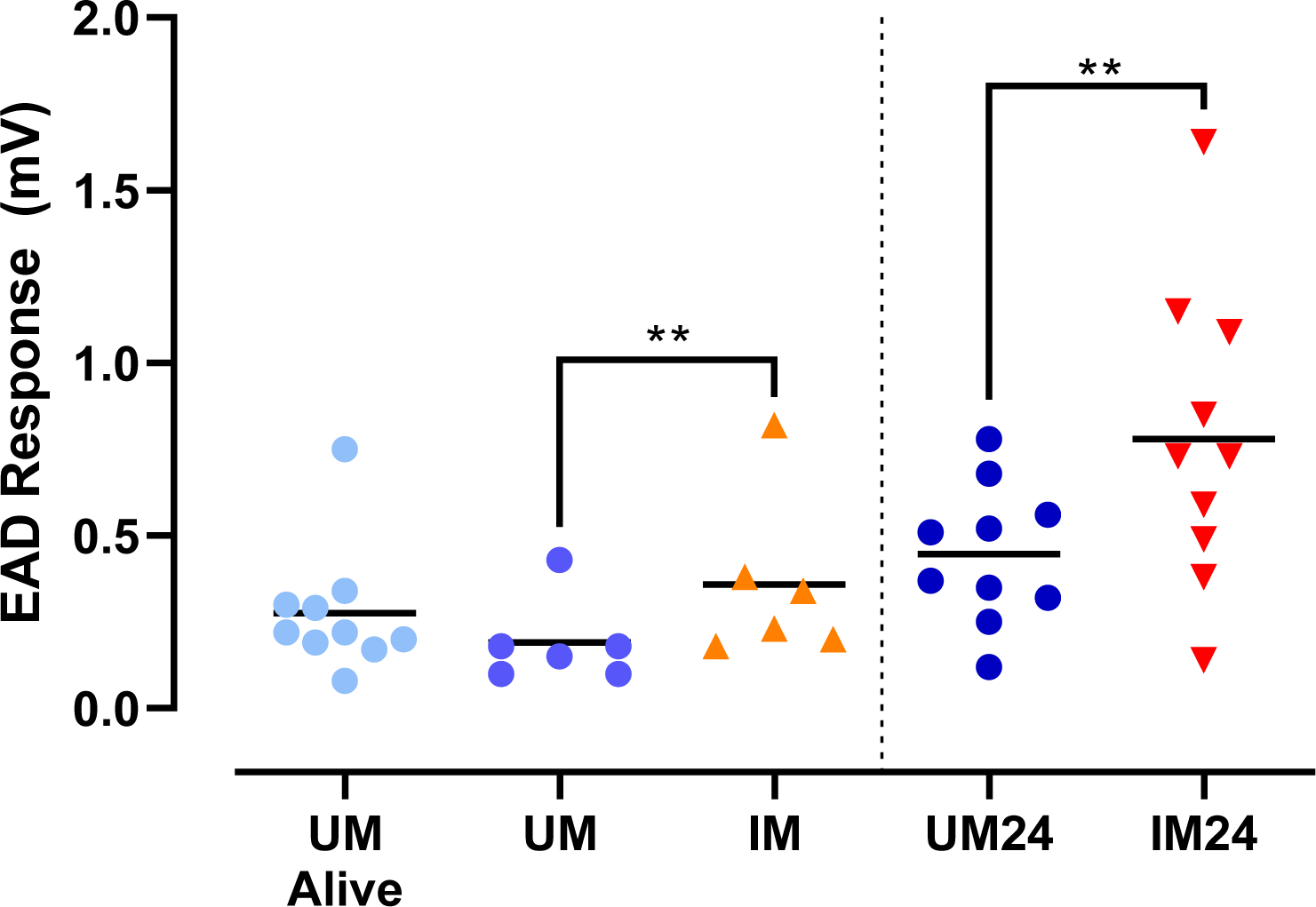
Male EAD response to volatiles from male cadavers. EAD responses (mV) to stimulus pipettes containing either uninfected, living males (UM Alive), uninfected male cadavers (UM), infected male cadavers (IM), Uninfected late-stage male cadavers (UM24), Infected late-stage male cadavers (IM24). There was a significant effect of treatment on the EAD response when comparing IM to UM (linear mixed-effects model (LMM), p < 0.01, t = 3.835) and when comparing IM24 to UM24 (LMM, p < 0.01, t = 3.887). (Significance reported as * p < 0.05; ** p < 0.01; *** p < 0.001).

**Supplementary figure 7.**
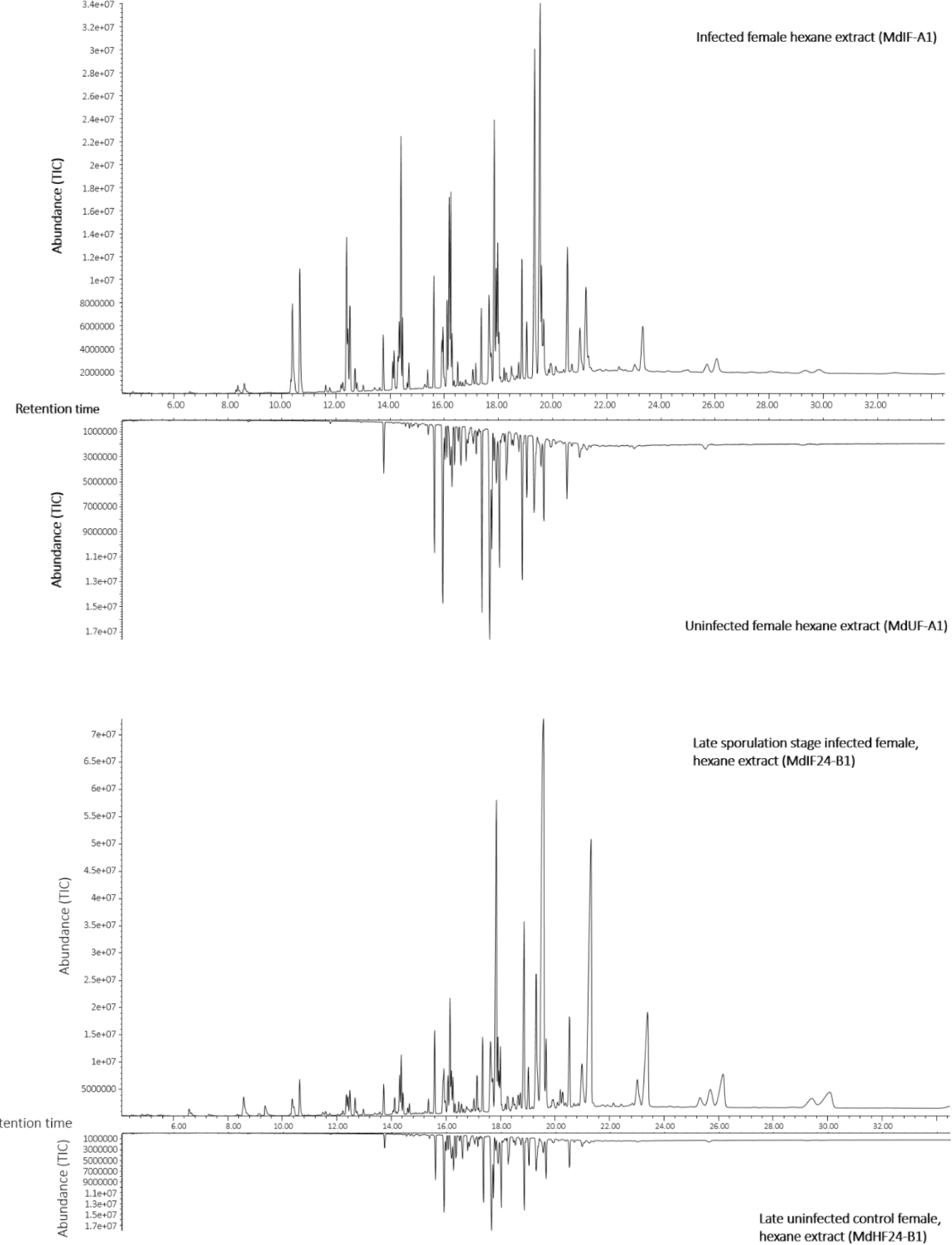
Gas chromatography mass spectrometry (GC-MS) representative Total Ion Chromatograms (TIC) of early and late sporulation stage infected female cadavers. GC-MS analysis conducted on cuticular hexane extracts of E. muscae early sporulation stage infected and uninfected female cadavers (top), and late sporulation stage infected and uninfected female control (bottom). Late killed cadavers had been incubated at high humidity for 26-28 hours, to prevent desiccation during sporulation.

**Supplementary figure 8.**
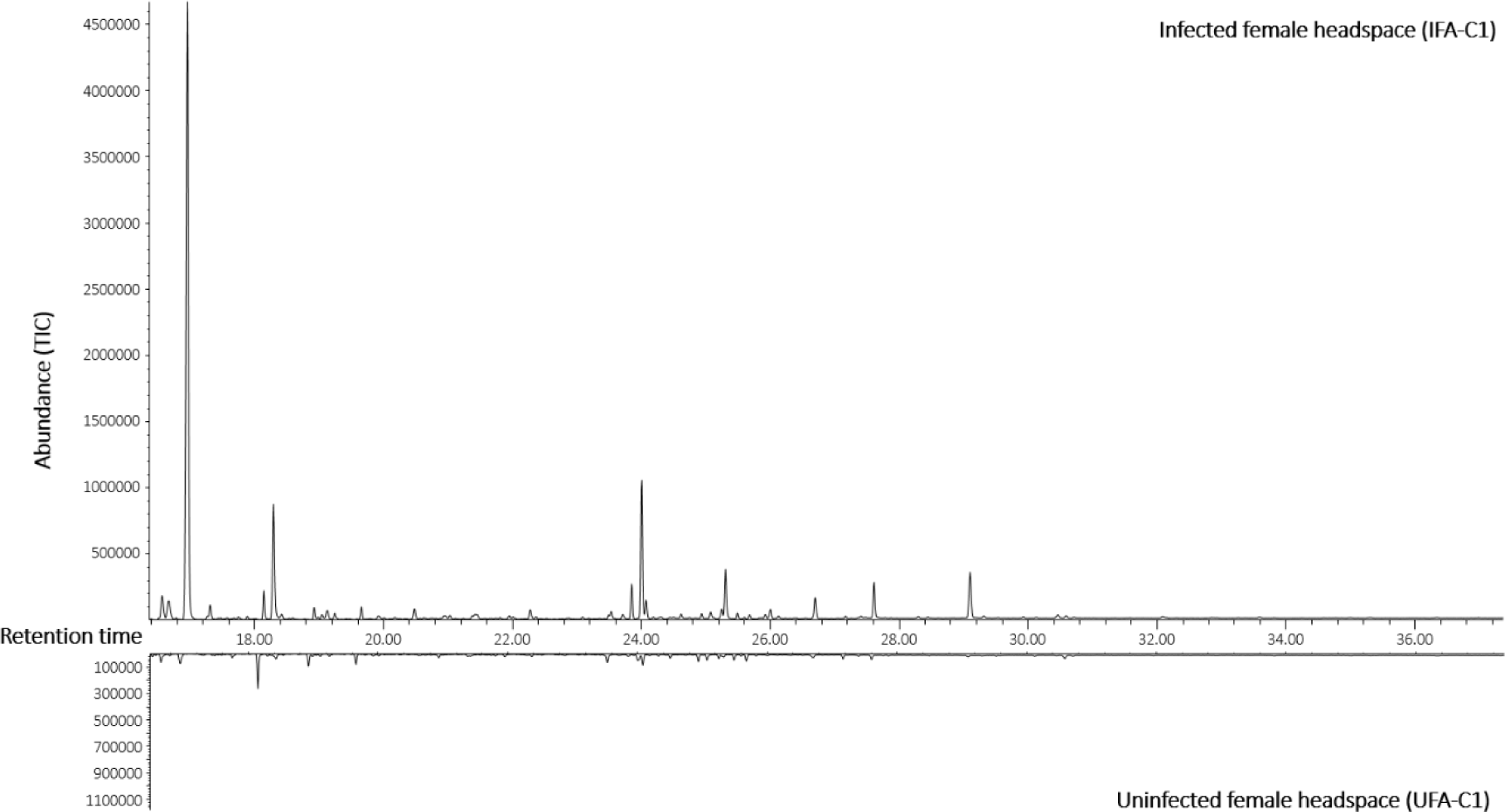
Gas chromatography mass spectrometry (GC-MS) representative Total Ion Chromatograms (TIC)of headspace samples from E. muscae sporulating females (top) and uninfected females (bottom). Chromatogram of headspace sampled as an aeration from female, sporulating houseflies (top). Headspace from uninfected female houseflies can be seen in the bottom.

**Supplementary figure 9.**
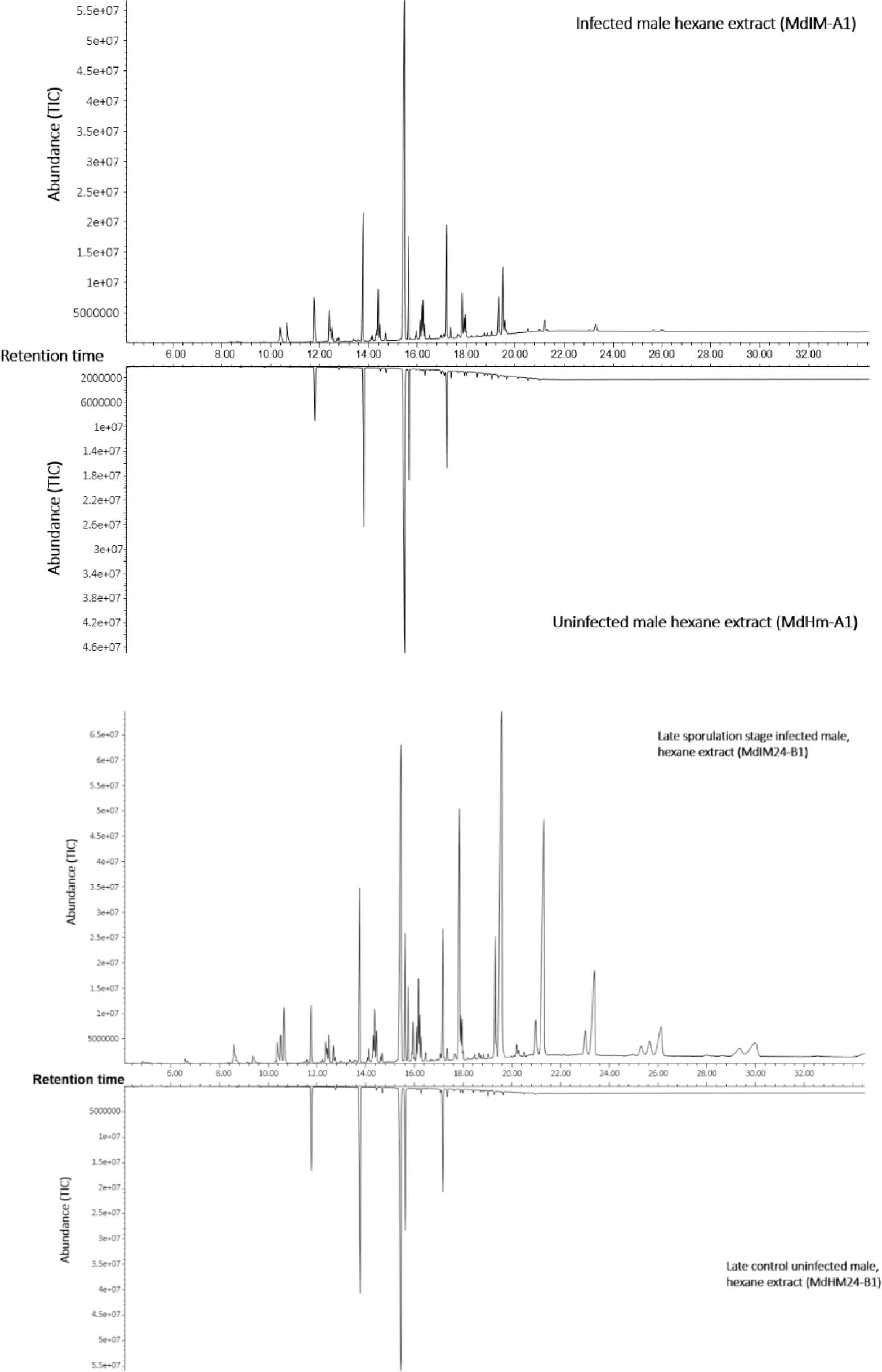
Representative Total Ion Chromatograms (TIC) of cuticular extacts of early (top) and late (bottom) sporulation stage infected male cadaver and an uninfected male control. Late killed cadavers had been incubated at high humidity for 24-26 hours, to prevent desiccation during sporulation.

**Supplementary figure 10.**
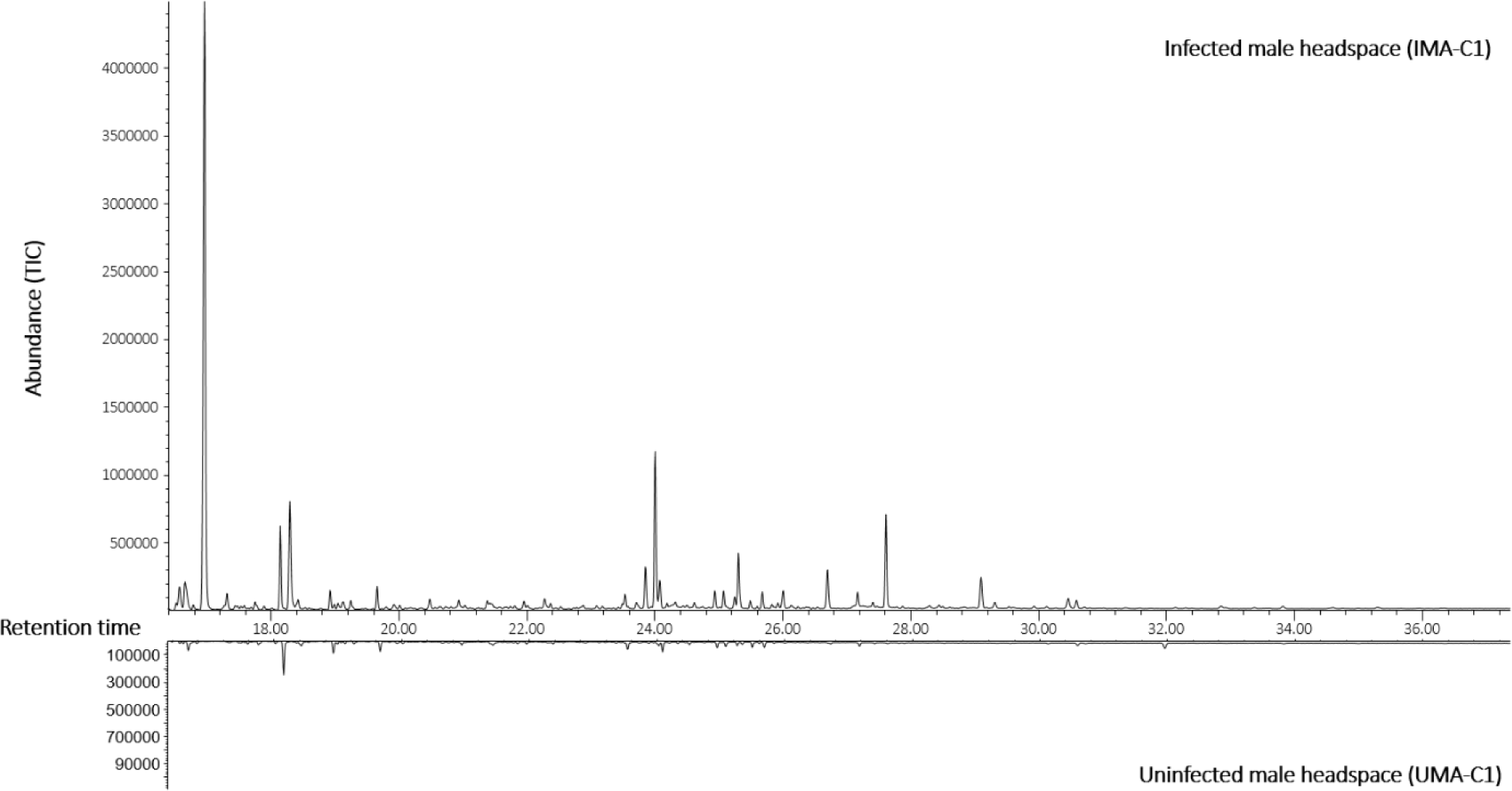
Total Ion Chromatogram (TOC) of headspace from male, sporulating houseflies (top) and from uninfected male houseflies (bottom).

**Supplementary figure 11.**
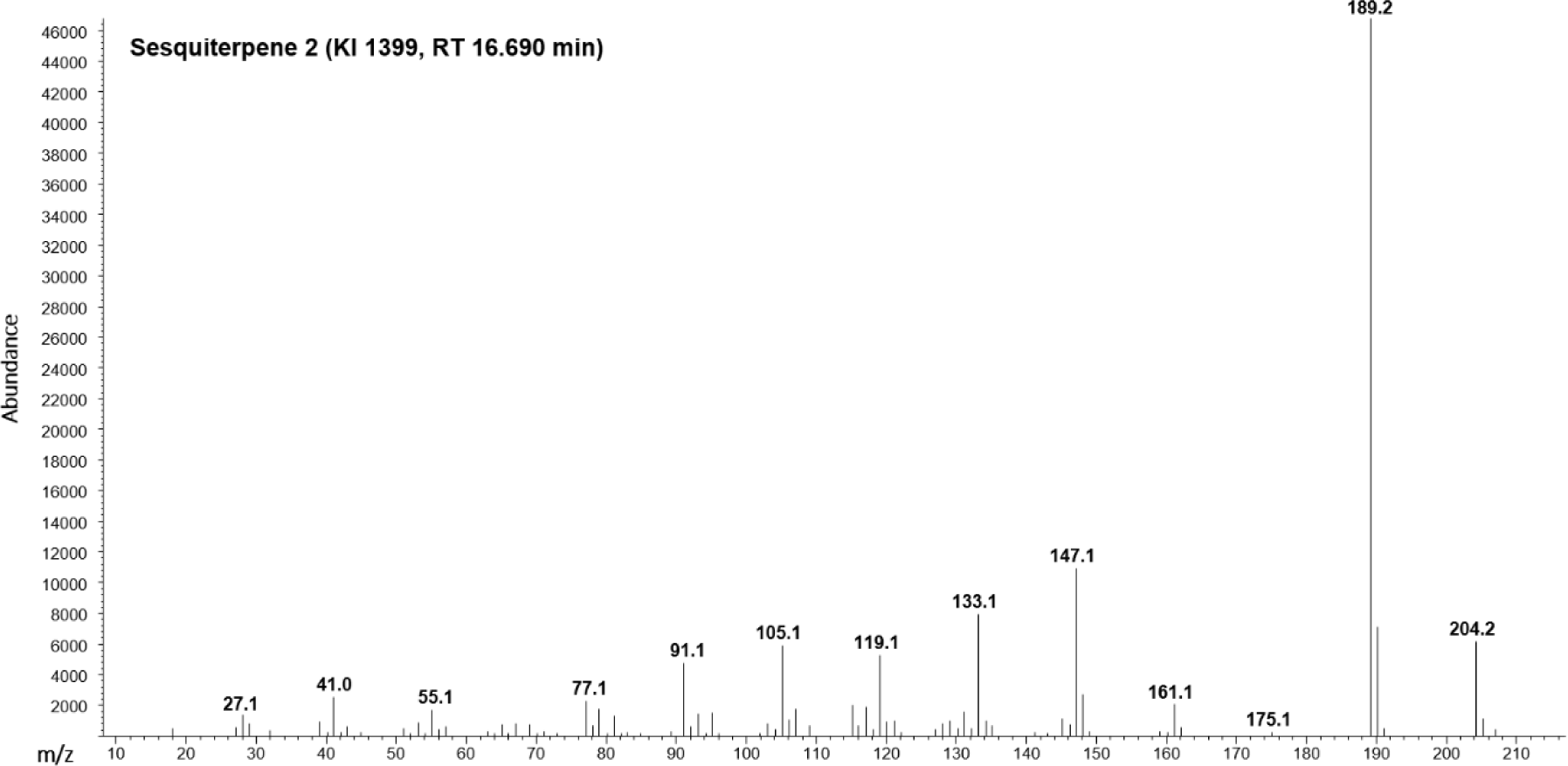
Mass spectrum of “Sesquiterpene 2”, one of the two unidentified GC-EAD active sesquiterpene compounds.

**Supplementary figure 12.**
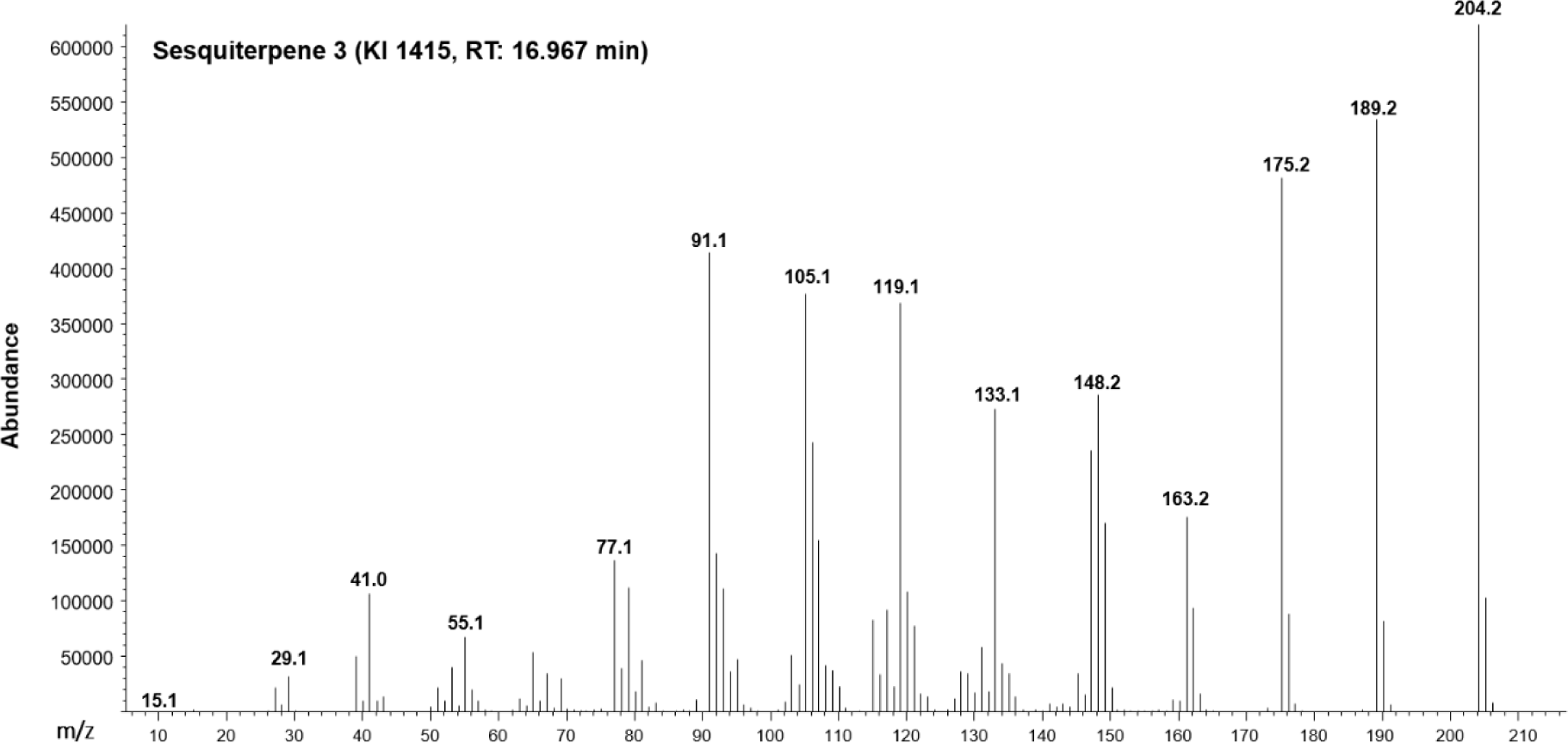
Mass spectrum of “Sesquiterpene 3”, the largest occurring compound in headspace from any *E. muscae* sample.

**Supplementary figure 13.**
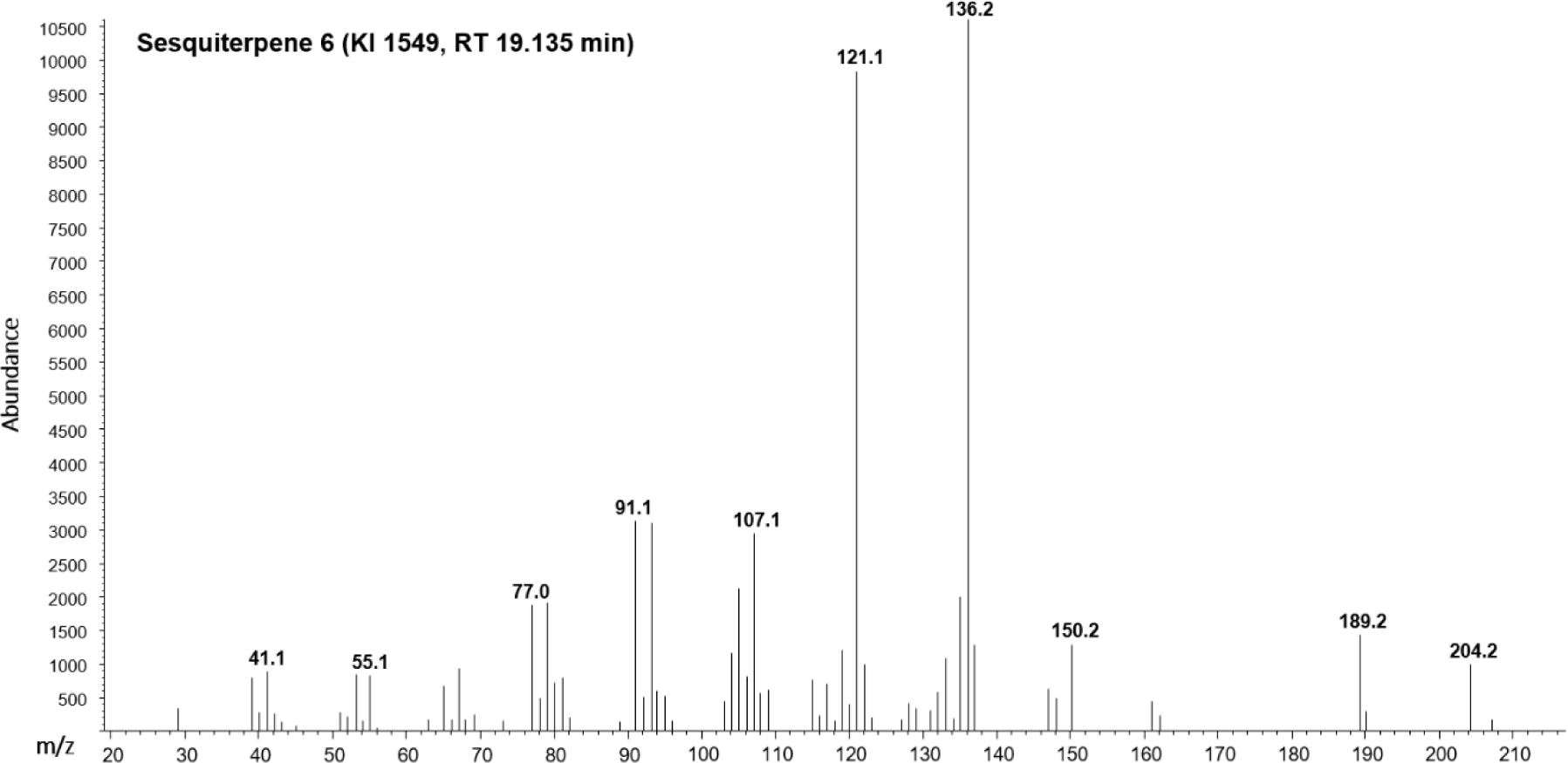
Mass spectrum of “Sesquiterpene 6”, the second GC-EAD active sesquiterpene.

**Supplementary figure 14.**
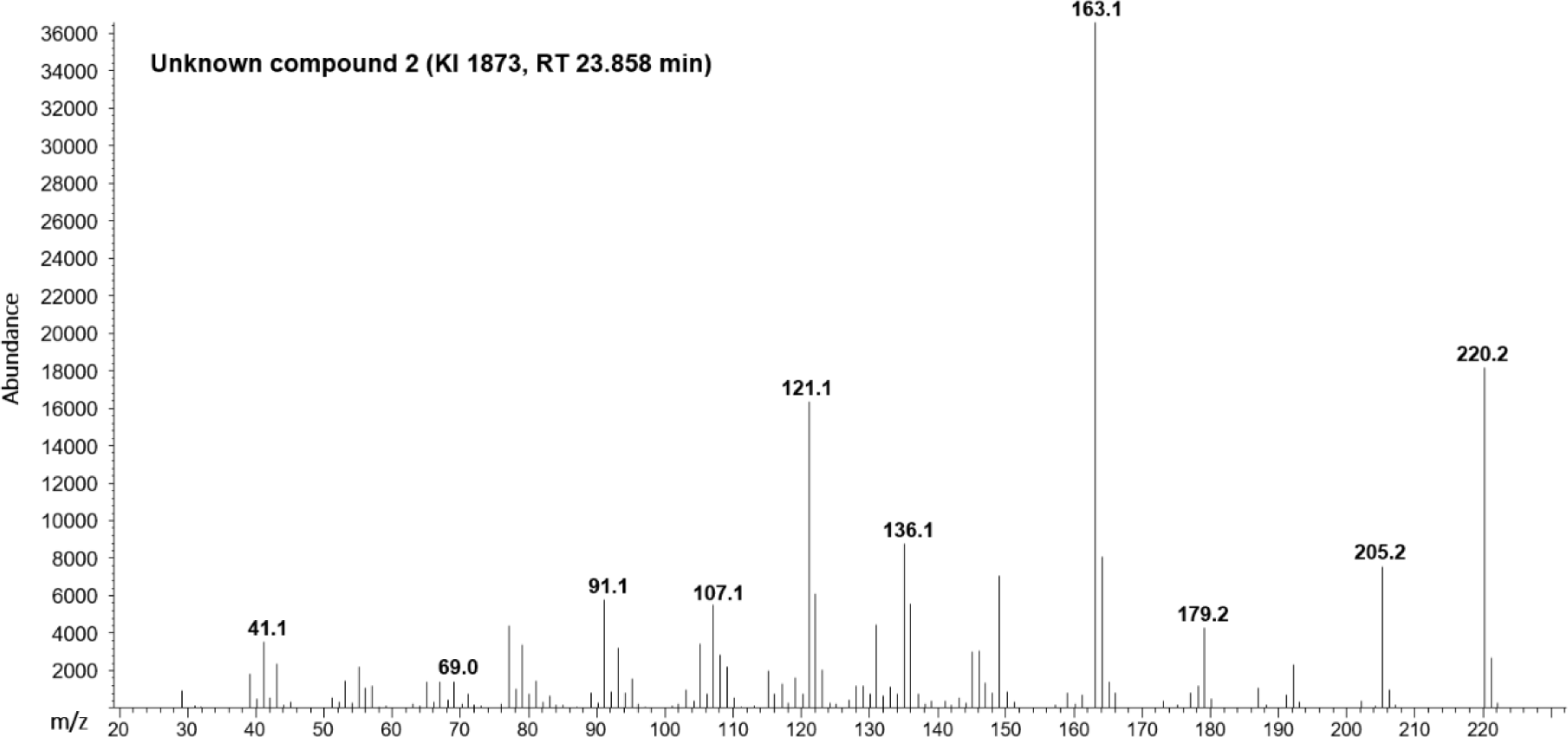
Mass spectrum of ”Unknown compound 2”, one of two GC-EAD compounds with unknown structure and identity.

**Supplementary figure 15.**
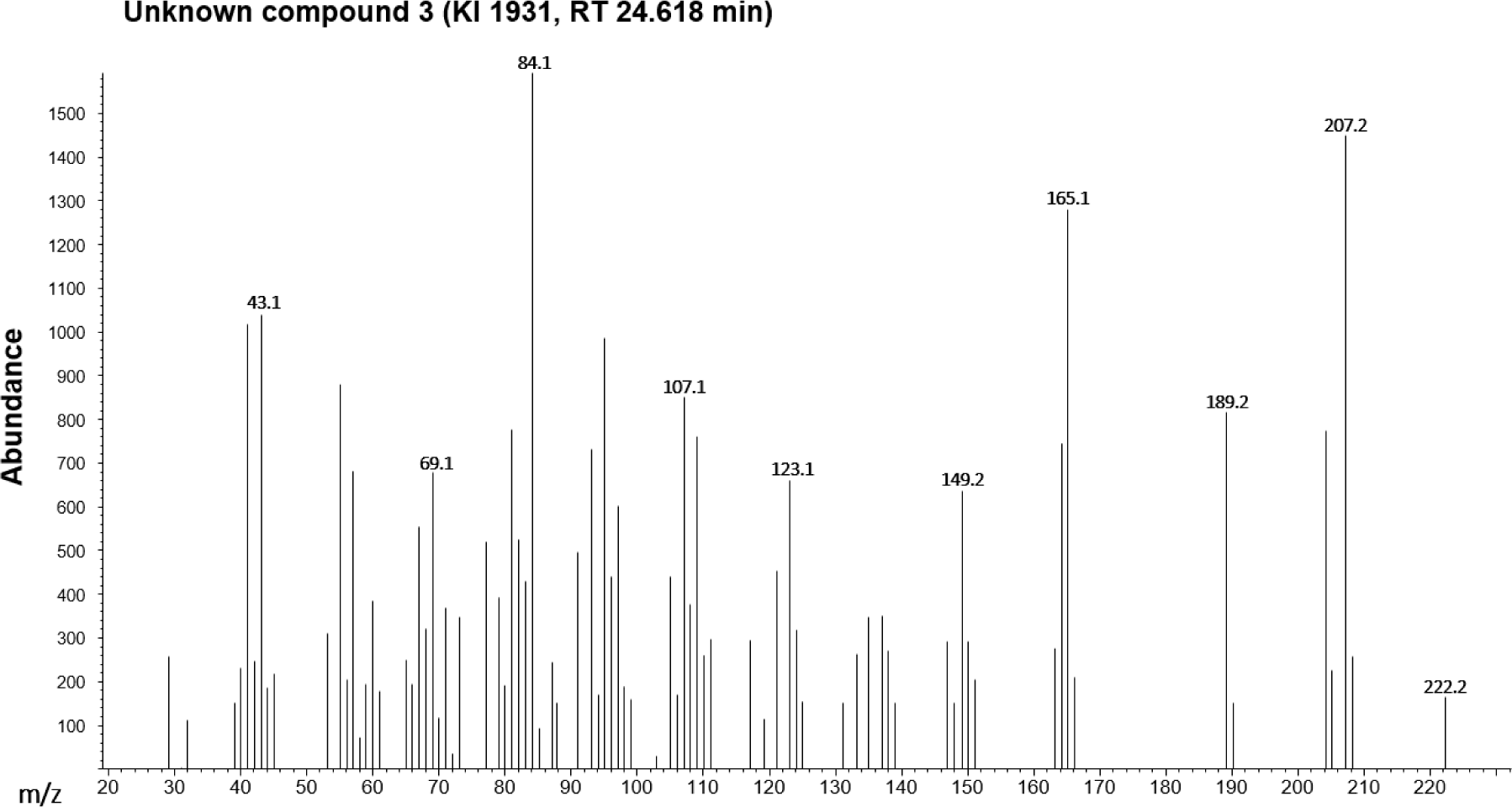
Mass spectrum of ”Unknown compound 3”, the second of two GC-EAD compounds with unknown structure and identity.

**Supplementary figure 16.**
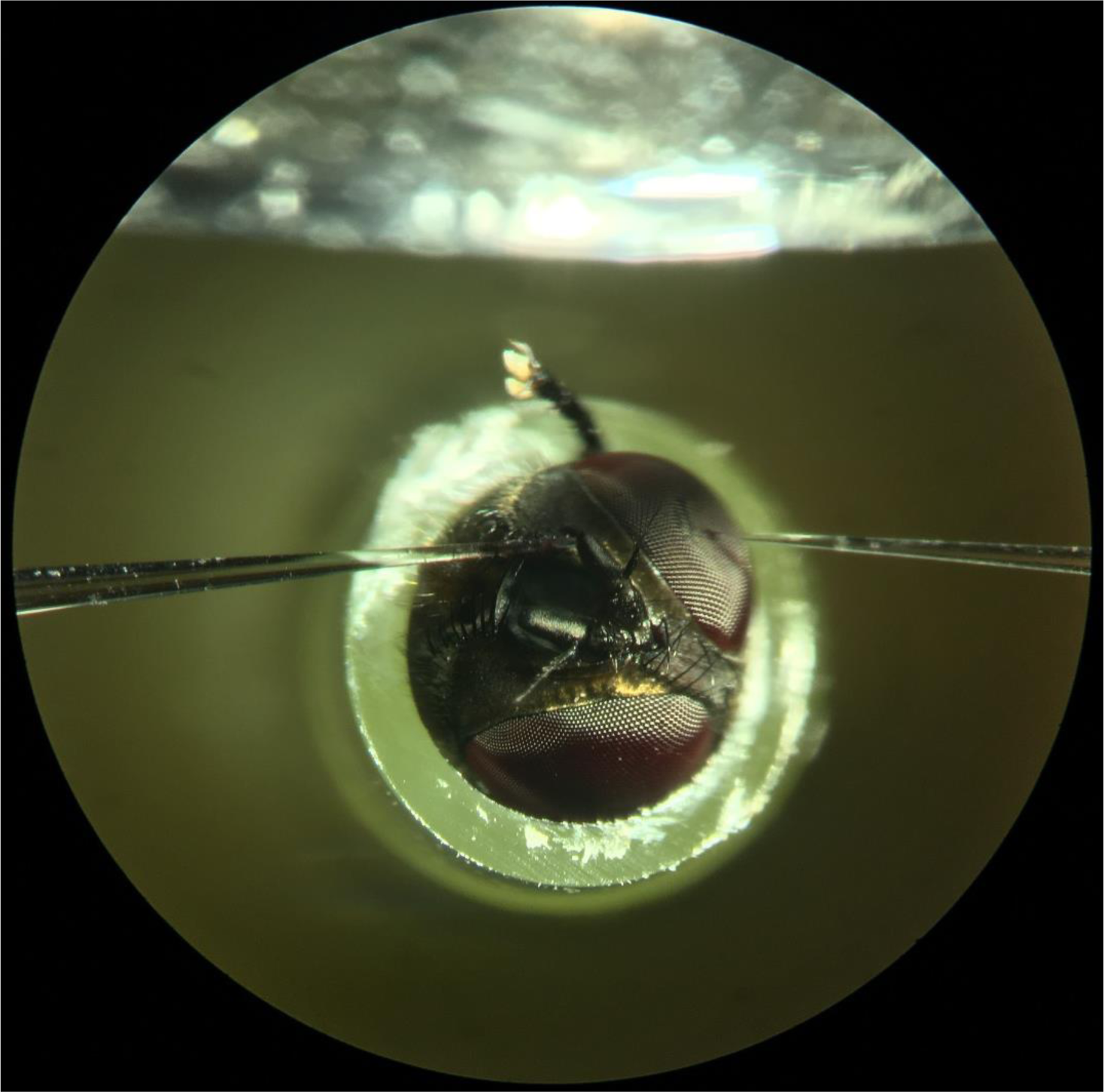
GC-EAD setup. A living, male housefly were fixated in a cut-off pipette tip so that only the head protruded. Electrodes in glass capillaries filled with Ringer solution were placed in eye (ground electrode) and on the tip of the funiculus (recording electrode) on the fly. The top, white glass surface is the glass tube that delivered purified and humidified air stream with compounds eluting from the the GC column.

**Supplementary figure 17.**
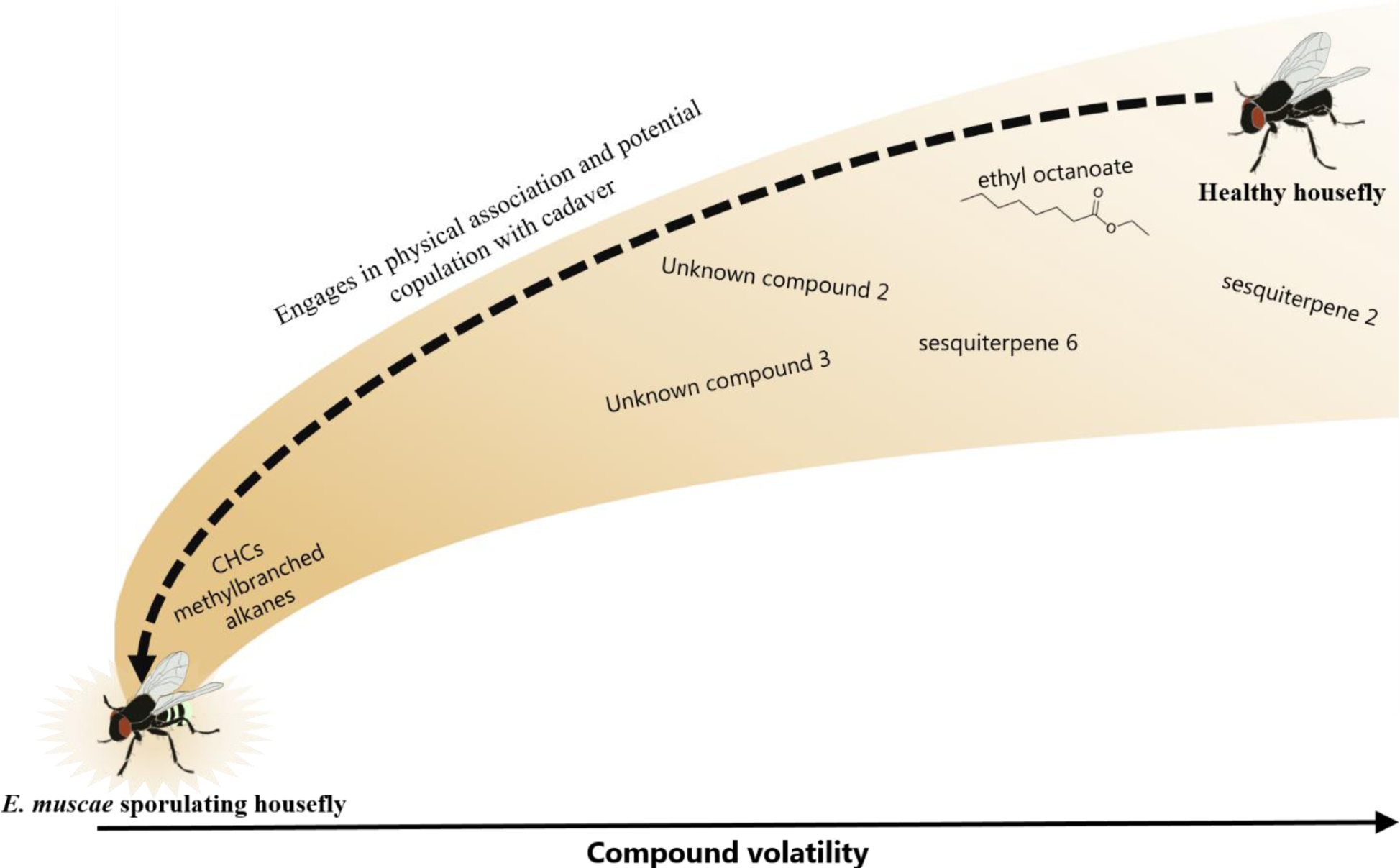
Conceptual model of housefly attraction to *E. muscae* sporulating cadavers. A healthy housefly male encounters a plume of highly volatile attractants. The male follows this increasing gradient of chemical cues until in vicinity of a sporulating female cadaver (dashed arrow). Here, the changes in the CHC profile and the methylbranched alkanes in particular, stimulate male mating behavior and cause him to engage in copulation with the cadaver.

**Supplementary data 1.** Percentage of male flies dying of *E. muscae* infection after exposure to sporulating cadavers. Males used in mating activity experiment were subsequently anaesthetized with CO_2_ and kept in individual cages with food and water ad libitum. All males were monitored for 10 days and assessed for successful *E. muscae* infection (shown by death and visible E. muscae conidiophores growing from the cadaver). No males exposed to freeze-killed cadavers (controls) developed *E. muscae* infections.

| Treatment | Percentage of males showing successful *E. muscae* infection |
| --- | --- |
| **M-IF** (Male to infected female, early sporulation stage) | 15.4 % (n= 13) |
| **M-IF24** (Male to infected female, late sporulation stage) | 73.33 % (n=15) |
| **M-IF24S** (Male to infected female, late sporulation stage with spore halo) | 68.42 % (n= 14) |
| **M-IM** (Male to infected male, early sporulation stage) | 9,09 % (n=11) |
| **M-IM24** (Male to infected male, late sporulation stage) | 18,18 % (n=11) |
| **M-IM24S** (Male to infected male, late sporulation stage with spore halo) | 82,35 % (n=17) |

**Supplementary data 2.**
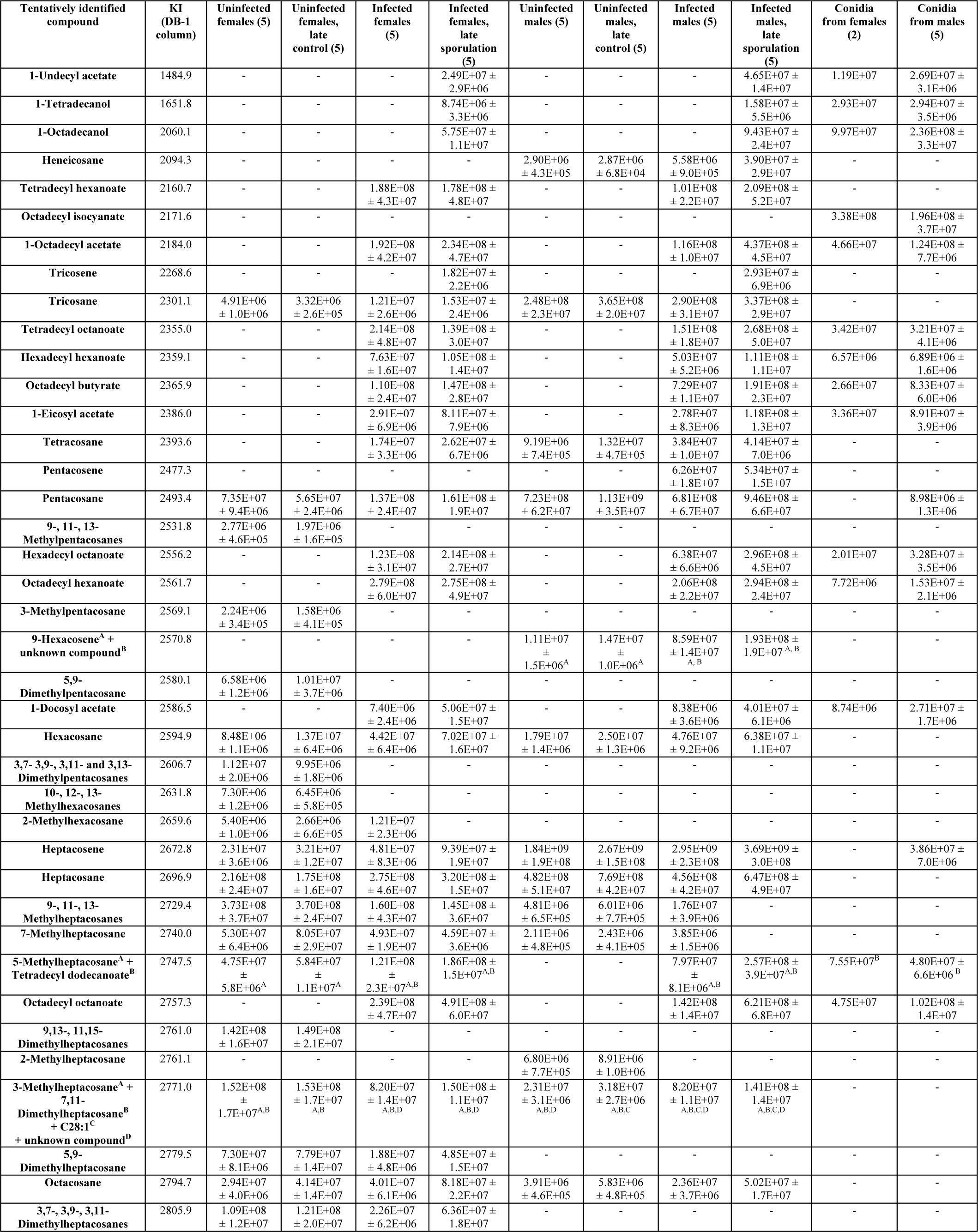
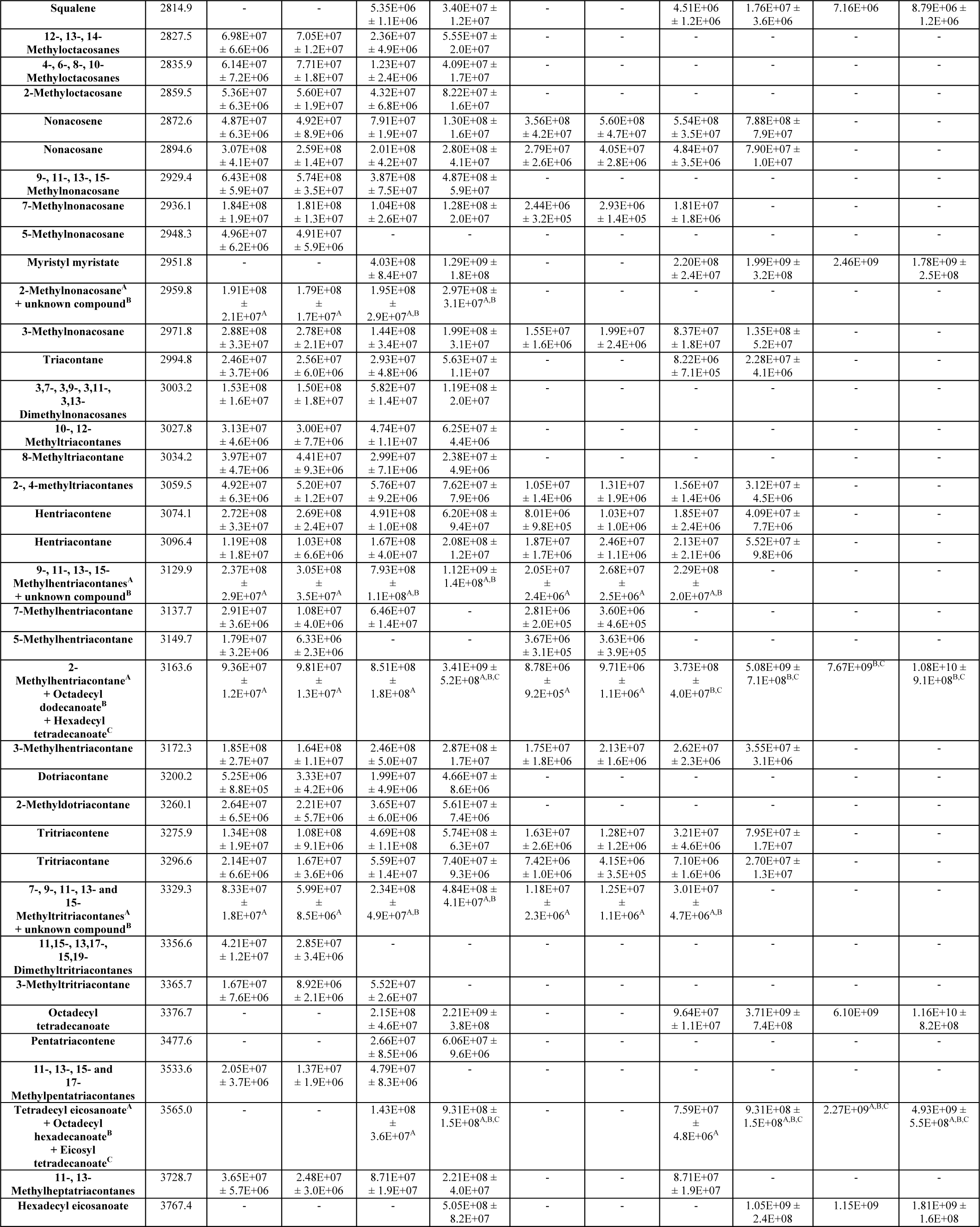
Compound table of housefly cuticular extracts. Full table of tentatively identified compounds in cuticular hexane extracts. First column shows compound names. Second columns shows Kovats Retention index. The relative amount of each compound in each sample type is given in peak area abundance ± Standard error of the mean. Parenthesis after each sample type is the number of samples. A compound had to appear in all 5 samples to be included. Unidentified compounds were not included.

**Supplementary data 3.** Full table of tentatively identified compounds in headspace samples. First column shows compound names. Second column shows Kovats Retention index. The relative amount of each compound in each sample type is given in peak area (TIC) ± Standard error of the mean. Parenthesis after each sample type is the number of samples. Compounds appearing in three out of five samples in a given treatment was included.

| Compound name | Kovats RI<br>(DB-WAX<br>column) | Uninfected females<br>(5) | Uninfected males<br>(5) | Sporulating females<br>(5) | Sporulating males<br>(5) | Conidia from females<br>(3) | Conidia from males<br>(2) |
| --- | --- | --- | --- | --- | --- | --- | --- |
| Sesquiterpene 1 | 1393 | - | - | 8.40E+06 ± 3.0E+06 | 8.56E+06 ± 3.1E+06 | - | - |
| Sesquiterpene 2 | 1399 | - | - | 4.99E+06 ± 2.2E+05 | 5.50E+06 ± 7.2E+05 | - | - |
| Sesquiterpene 3 | 1415 | 3.76E+06 ± 1.5E+06 | - | 1.46E+08 ± 6.2E+07 | 1.67E+08 ± 5.6E+07 | 6.75E+06 ± 1.3E06 | 5.73E+06 |
| Ethyl octanoate | 1436 | - | - | 2.81E+07 ± 2.0E+07 | 5.76E+06 ± 2.2E+06 | - | - |
| Sesquiterpene 4 | 1472 | - | - | 1.31E+06 ± 6.0E+05 | 1.27E+06 ± 5.5E+05 | - | - |
| Sesquiterpene 5 | 1496 | - | - | 1.31E+07 ± 4.0E+06 | 3.56E+07 ± 1.6E+07 | - | - |
| Sesquiterpene 6 | 1549 | - | - | 1.89E+06 ± 8.1E+05 | 2.38E+06 ± 1.1E+06 | - | - |
| Sesquiterpene 7 | 1636 | - | - | 4.44E+06 ± 1.7E+06 | 2.97E+06 ± 1.2E+06 | - | - |
| Sesquiterpene 8 | 1673 | - | - | 3.55E+05 ± 5.0E+04 | 3.95E+05 ± 5.5E+04 | - | - |
| Unknown compound 1 | 1758 | - | - | 5.04E+06 ± 3.4E+06 | 4.68E+06 ± 2.8E+06 | - | - |
| Unknown compound 2 | 1873 | - | - | 1.44E+07 ± 6.9E+06 | 1.52E+07 ± 6.4E+06 | - | - |
| Unknown | 1886 | - | - | 1.79E+07 ± 4.0E+06 | 5.19E+07 ± 1.9E+07 | - | - |
| 1-Phenylethyl alcohol | 1922 | - | 5.17E+06 ± 3.1E+06 | - | - | - | - |
| Unknown compound 3 | 1931 | - | - | 4.73E+05 ± 7.0E+04 | 3.72E+06 ± 2.0E+06 | - | - |
| Unknown compound 4 | 1980 | - | - | 5.38E+06 ± 2.6E+06 | 1.26E+06 ± 2.5E+05 | - | - |
| Unknown compound 5 | 1984 | - | - | 1.85E+07 ± 9.5E+06 | 2.01E+07 ± 9.1E+06 | - | - |
| Unknown compound 6 | 2033 | - | - | 2.55E+06 ± 1.4E+06 | 5.39E+05 ± 1.0E+05 | - | - |
| Unknown compound 7 | 2039 | - | - | 6.69E+06 ± 3.0E+06 | 6.82E+06 ± 2.6E+06 | - | - |
| Unknown compound 8 | 2050 | - | - | 8.79E+05 ± 3.2E+05 | 4.51E+05 ± 1.3E+05 | - | - |
| 1-Tetradecyl acetate | 2094 | - | - | 1.85E+07 ± 1.1E+07 | 3.68E+07 ± 1.9E+07 | 1.10E+07 ± 2.9E+06 | 1.52E+07 |
| 1-Tetradecanol | 2168 | 3.91E+06 ± 2.1E+06 | 2.53E+06 ± 1.3E+06 | 2.33E+07 ± 1.2E+07 | 2.49E+07 ± 1.2E+07 | 8.07E+06 ± 1.5+06 | 1.15E+07 |
| Unknown compound 9 | 2226 | - | - | 3.90E+05 ± 3.0E+04 | 4.25E+05 ± 6.7E+04 | - | - |
| Diterpene 1 | 2286 | - | - | 2.75E+06 ± 7.9E+05 | 2.66E+06 ± 8.6E+05 | - | - |
| 1-Hexadecyl acetate | 2300 | - | - | 9.10E+06 ± 6.3E+06 | 2.26E+06 ± 1.1E+06 | - | - |
| 1-Hexadecanol | 2376 | - | - | 3.79E+06 ± 1.8E+06 | 4.30E+06 ± 1.7E+06 | - | - |

**Supplementary data 4.** Fatty acid gene expression in *E. muscae*. **A**. Schematic drawing of the general fatty-acid synthesis pathway in fungi. **B**. Expressed *E. muscae* transcripts annotated as either Acetyl-CoA carboxylase, Fatty acid synthases, or long-chain acyl-CoA synthetase enzymes. Fungal transcripts written in bold are significantly higher expressed in late-stage sporulating cadavers vs. early-stage sporulating cadavers.

|  | TranscriptID | BLASTPtophit | %Identity | e-value | Kegg | EC | log2FC | FDR_p |
| --- | --- | --- | --- | --- | --- | --- | --- | --- |
| Acetyl-CoA carboxylase | TRINITY_DN20838_c2_g2 | ACAC_SCHPO | 39,3 | 8.14e-95 | K11262 | 6.4.1.2, 6.3.4.14 | 2,46 | 0,0000 |
|  | TRINITY_DN9224_c0_g1 | ACAC_SCHPO | 66,7 | 6.98e-19 | K11262 | 6.4.1.2, 6.3.4.14 | 4,19 | 0,0000 |
|  | TRINITY_DN11999_c0_g1 | ACAC_SCHPO | 37,7 | 2.04e-61 | K11262 | 6.4.1.2, 6.3.4.14 | 2,66 | 0,0000 |
|  | TRINITY_DN8745_c0_g1 | ACAC_YEAST | 70,3 | 1.59e-117 | K11262 | 6.4.1.2, 6.3.4.14 | 3,46 | 0,0001 |
|  | TRINITY_DN1774_c0_g1 | ACAC_YEAST | 44,0 | 5.42e-49 | K11262 | 6.4.1.2, 6.3.4.14 | 2,56 | 0,0002 |
|  | TRINITY_DN6921_c0_g1 | ACAC_SCHPO | 62,7 | 2.77e-41 | K11262 | 6.4.1.2, 6.3.4.14 | 4,06 | 0,0010 |
|  | TRINITY_DN5496_c0_g1 | ACAC_YEAST | 54,6 | 3.58e-62 | K11262 | 6.4.1.2, 6.3.4.14 | 3,76 | 0,0024 |
|  | TRINITY_DN10159_c0_g1 | ACAC_SCHPO | 58,6 | 3.63e-49 | K11262 | 6.4.1.2, 6.3.4.14 | 1,38 | 0,0035 |
|  | TRINITY_DN20838_c1_g1 | ACAC_SCHPO | 31,7 | 2.92e-15 | K11262 | 6.4.1.2, 6.3.4.14 | -0,76 | 0,0047 |
|  | TRINITY_DN21736_c0_g1 | ACAC_SCHPO | 62,1 | 3.35e-70 | K11262 | 6.4.1.2, 6.3.4.14 | 1,69 | 0,0059 |
|  | TRINITY_DN1062_c0_g1 | ACAC_SCHPO | 53,3 | 4.62e-40 | K11262 | 6.4.1.2, 6.3.4.14 | 1,50 | 0,0174 |
|  | TRINITY_DN21389_c0_g1 | ACAC_SCHPO | 24,1 | 2.56e-22 | K11262 | 6.4.1.2, 6.3.4.14 | 0,47 | 0,0195 |
|  | TRINITY_DN20838_c2_g1 | ACAC_SCHPO | 50,6 | 0 | K11262 | 6.4.1.2, 6.3.4.14 | 0,39 | 0,0568 |
|  | TRINITY_DN2573_c0_g1 | ACAC_SCHPO | 61,3 | 5.9e-28 | K11262 | 6.4.1.2, 6.3.4.14 | 1,78 | 0,0591 |
|  | TRINITY_DN2535_c0_g1 | ACAC_YEAST | 79,2 | 6.31e-64 | K11262 | 6.4.1.2, 6.3.4.14 | 2,31 | 0,0741 |
|  | TRINITY_DN5348_c0_g1 | ACAC_YEAST | 75,4 | 5.07e-66 | K11262 | 6.4.1.2, 6.3.4.14 | 2,37 | 0,0925 |
| Fatty acid synthases FAS1/FAS2 | TRINITY_DN21615_c2_g1 | FAS2_PENPA | 54,4 | 0 | K00667 | 2.3.1.86 | 2,45 | 0,0000 |
|  | TRINITY_DN20289_c0_g1 | FAS2_EMEND | 52,5 | 0 | K00667 | 2.3.1.86 | 3,43 | 0,0000 |
|  | TRINITY_DN21615_c2_g3 | FAS2_YEAST | 65,3 | 6.1e-36 | K00667 | 2.3.1.86 | 3,13 | 0,0000 |
|  | TRINITY_DN16765_c0_g1 | FAS2_EMEND | 64,4 | 1.62e-133 | K00667 | 2.3.1.86 | 2,25 | 0,0000 |
|  | TRINITY_DN21615_c0_g1 | FAS2_LACKL | 53,6 | 2.14e-35 | K00667 | 2.3.1.86 | 3,07 | 0,0014 |
|  | TRINITY_DN7801_c0_g1 | FAS2_CANAX | 81,0 | 1.63e-39 | K00667 | 2.3.1.86 | 1,59 | 0,0017 |
|  | TRINITY_DN1128_c0_g1 | FAS2_PENPA | 76,1 | 2.49e-32 | K00667 | 2.3.1.86 | 3,30 | 0,0154 |
|  | TRINITY_DN2844_c0_g1 | FAS2_SCHPO | 69,6 | 9.34e-32 | K00667 | 2.3.1.86 | 2,24 | 0,1581 |
|  | TRINITY_DN21615_c2_g4 | FAS1_SCHPO | 49,4 | 0 | K00668 | 2.3.1.86 | 3,33 | 0,0000 |
|  | TRINITY_DN17570_c0_g1 | FAS1_SCHPO | 58,1 | 9.62e-149 | K00668 | 2.3.1.86 | 3,99 | 0,0000 |
|  | TRINITY_DN12197_c0_g1 | FAS1_YEAST | 49,3 | 4.23e-83 | K00668 | 2.3.1.86 | 3,75 | 0,0000 |
|  | TRINITY_DN17406_c0_g1 | FAS1_SCHPO | 51,5 | 4.78e-115 | K00668 | 2.3.1.86 | 4,82 | 0,0000 |
|  | TRINITY_DN15402_c0_g1 | FAS1_YARLI | 39,8 | 1.57e-61 | K00668 | 2.3.1.86 | 4,65 | 0,0000 |
|  | TRINITY_DN14812_c0_g1 | FAS1_CANAX | 36,2 | 1.26e-37 | K00668 | 2.3.1.86 | 3,98 | 0,0002 |
|  | TRINITY_DN3608_c0_g1 | FAS1_YEAST | 50,7 | 3.45e-20 | K00668 | 2.3.1.86 | 2,57 | 0,0831 |
|  | TRINITY_DN5417_c0_g1 | FAS1_SCHPO | 63,6 | 4.8e-37 | K00668 | 2.3.1.86 | 2,93 | 0,1534 |
| long-chain acyl-CoA synthetase | TRINITY_DN17778_c0_g2 | LCF2_YEAST | 31,1 | 3.06e-49 | K01897 | 6.2.1.3 | 1,49 | 0,0000 |
|  | TRINITY_DN25342_c0_g1 | LCF2_SCHPO | 34,8 | 1.13e-136 | K01897 | 6.2.1.3 | 1,70 | 0,0000 |
|  | TRINITY_DN19153_c0_g2 | LCF2_SCHPO | 38,3 | 4.99e-164 | K01897 | 6.2.1.3 | 0,87 | 0,0000 |
|  | TRINITY_DN17671_c0_g1 | LCF2_YEAST | 25,7 | 1.99e-43 | K01897 | 6.2.1.3 | 0,54 | 0,0000 |
|  | TRINITY_DN17778_c0_g1 | LCF2_YEAST | 35,6 | 2.46e-39 | K01897 | 6.2.1.3 | -0,50 | 0,0002 |
|  | TRINITY_DN19147_c0_g1 | LCF2_YEAST | 35,6 | 3.89e-110 | K01897 | 6.2.1.3 | -0,28 | 0,0004 |
|  | TRINITY_DN20446_c3_g2 | LCF2_SCHPO | 33,3 | 1.76e-121 | K01897 | 6.2.1.3 | 4,30 | 0,0008 |
|  | TRINITY_DN21065_c0_g2 | LCF2_YEAST | 31,4 | 3.05e-95 | K01897 | 6.2.1.3 | 0,61 | 0,0047 |
|  | TRINITY_DN20446_c3_g1 | LCF1_YEAST | 37,1 | 6,00E-106 | K01897 | 6.2.1.3 | 4,21 | 0,2630 |
|  | TRINITY_DN19131_c0_g1 | LCF2_YEAST | 33,4 | 5.81e-88 | K01897 | 6.2.1.3 | -0,12 | 0,5653 |

**Supplementary data 5.**
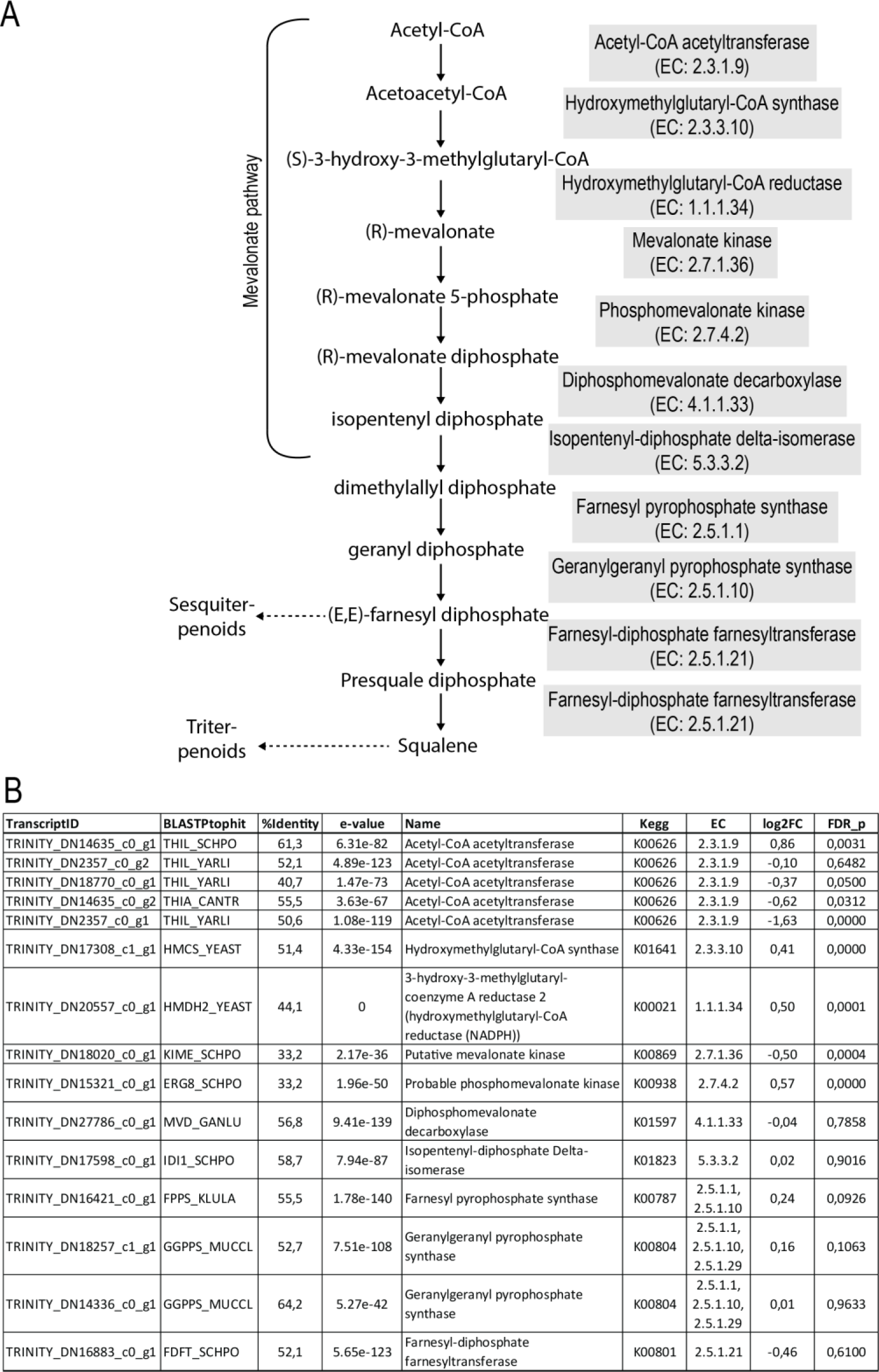
Mevalonate and terpenoid synthesis gene expression in *E. muscae*. **A**. Schematic drawing of the general terpenoid synthesis pathway in fungi. **B**. Expressed *E. muscae* transcripts annotated as enzymes in the general terpenoid synthesis pathway in fungi. Annotation, blastp results, and expression in late stage sporulating cadavers vs. early stage cadavers are given.

**Supplementary data 6.** Expressed *E. muscae* transcripts with homology to ethyl ester biosynthesis genes. **A**. Four E. muscae transcripts with homology the yeast *Saccharomyces cerevisiae* ethyl ester biosynthesis genes eht1 and eeb1 specifically involved in ethyl octanoate biosynthesis. Results of Blastp searches and expression in late-stage sporulating cadavers vs. early-stage sporulating cadavers are shown. **A.** Results of Blastp searches with the yeast ethyl ester biosynthesis genes eht1, eeb1, and mgl2 against the *E. muscae* translated transcripts

| TranscriptID | BLASTPtophit | Query/Hit_coverage | %Identity | e-value | Name | Kegg | EC | log2FC | FDR_p |
| --- | --- | --- | --- | --- | --- | --- | --- | --- | --- |
| TRINITY_DN19029_c0_g1 | PANB_EMENI | Q:736-1566,H:62-340 | 51,613 | 3.02e-84 | 3-methyl-2-oxobutanoate | K00606 | 2.1.2.11 | -0,38 | 0,0617 |
| **TRINITY_DN8643_c0_g1** | **YM60_YEAST** | **Q:925-362,H:207-397** | **27,586** | **1.36e-16** | **Putative esterase** | **K07019** | - | **2,96** | **0,0000** |
| TRINITY_DN19826_c0_g1 | YM60_YEAST | Q:1519-410,H:8-413 | 29,71 | 2.82e-52 | Putative esterase | K07019 | - | 0,42 | 0,0007 |
| TRINITY_DN18612_c0_g1 | YM60_YEAST | Q:314-1420,H:33-413 | 29,095 | 1.32e-42 | Putative esterase | K07019 | - | 0,21 | 0,2334 |

B
| Query = *S. cerevisiae eeb1* (GU471249.1) |  |  |
| --- | --- | --- |
| TranscriptID | Score (bits) | e-value |
| TRINITY_DN18612_c0_g1 | 102,0 | 2,00E-38 |
| TRINITY_DN19826_c0_g1 | 91,8 | 1,00E-33 |
| TRINITY_DN8643_c0_g1 | 46,4 | 9,00E-12 |

| Query = *S. cerevisiae mgl2* (YMR210W) |  |  |
| --- | --- | --- |
| TranscriptID | Score (bits) | e-value |
| TRINITY_DN19826_c0_g1 | 106,0 | 3,00E-46 |
| TRINITY_DN18612_c0_g1 | 105,0 | 3,00E-40 |
| TRINITY_DN8643_c0_g1 | 52,4 | 2,00E-13 |

| Query = *S. cerevisiae eht1* (AB012577.1) |  |  |
| --- | --- | --- |
| TranscriptID | Score (bits) | e-value |
| TRINITY_DN19826_c0_g1 | 96,4 | 3,00E-30 |
| TRINITY_DN18612_c0_g1 | 97,3 | 6,00E-29 |
| TRINITY_DN19029_c0_g1 | 88,1 | 3,00E-17 |
| TRINITY_DN8643_c0_g1 | 49,6 | 5,00E-07 |

**Supplementary data 7.**
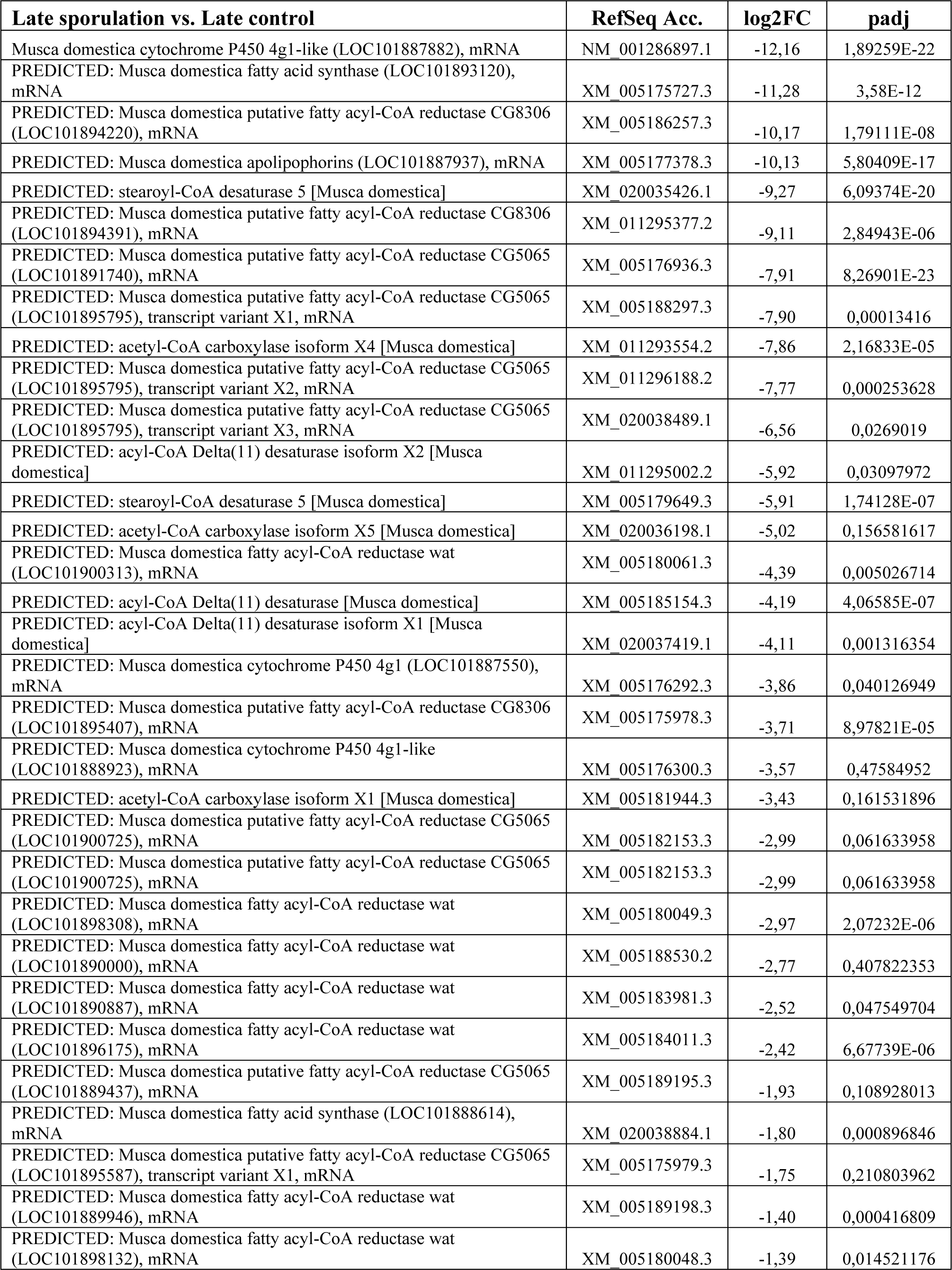
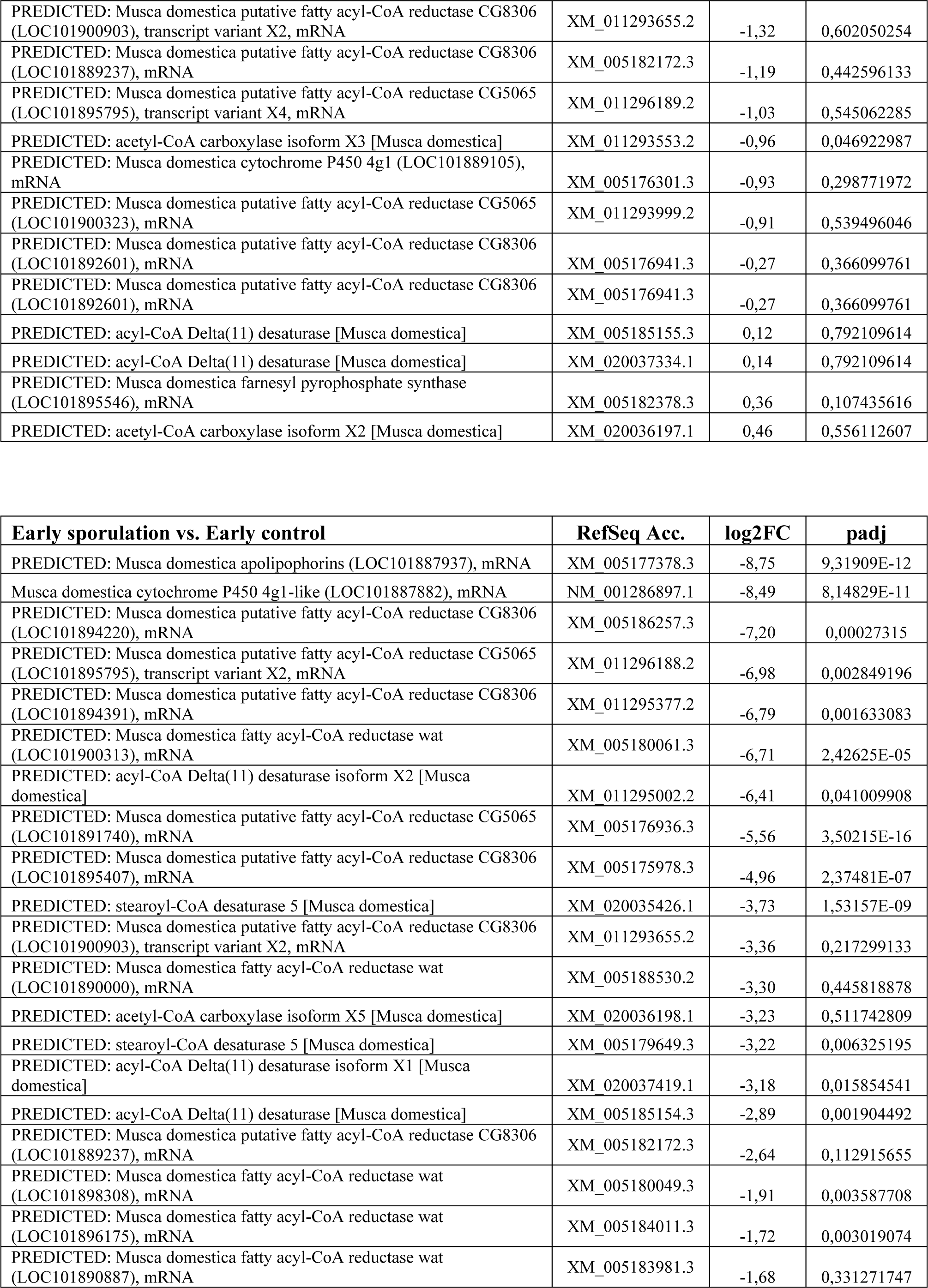
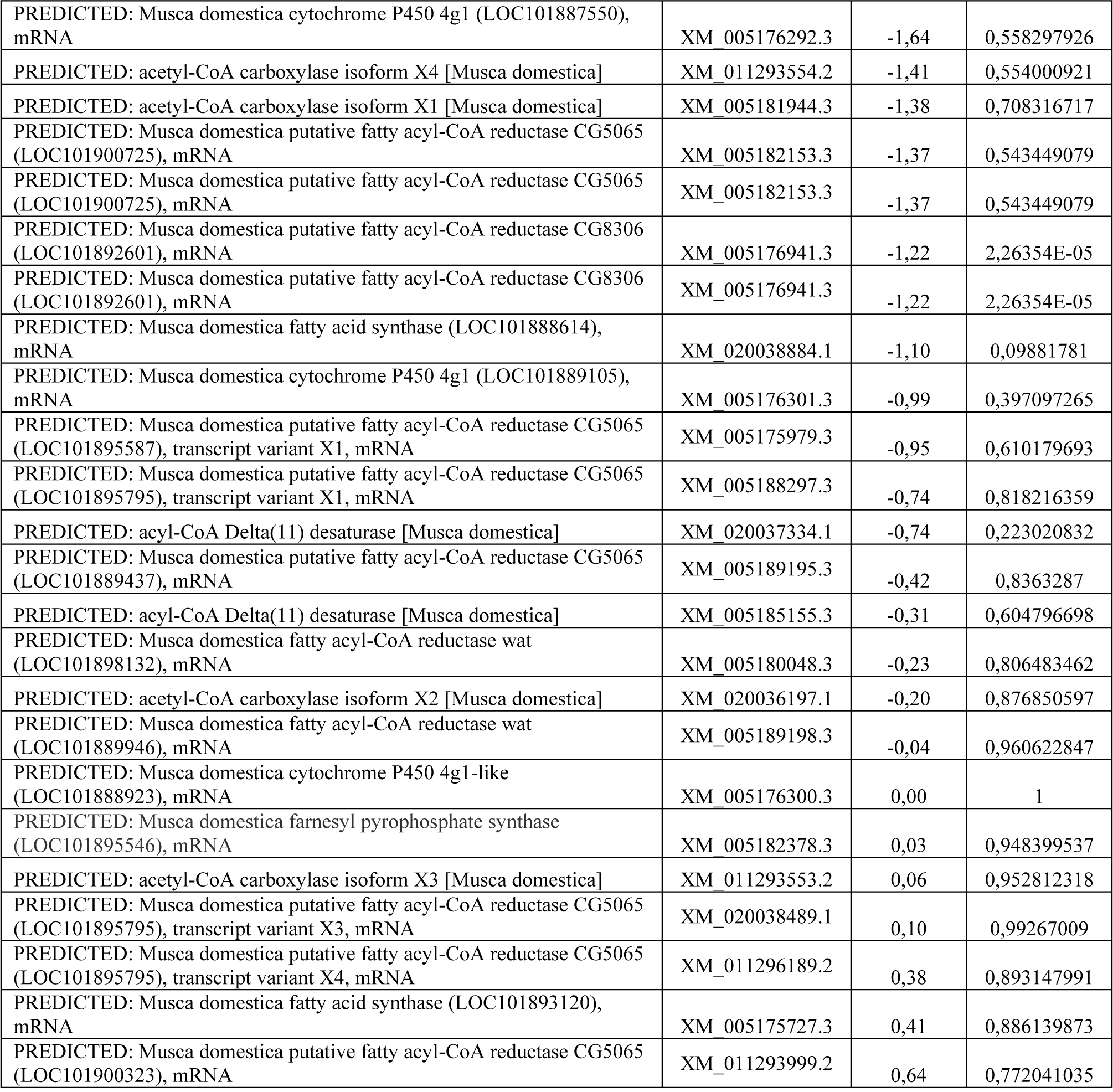
Housefly gene expression of selected cuticular hydrocarbon biosynthesis related genes. Tables are sorted by log2 Fold Change. Top table: Late *E. muscae* sporulating female house flies compared to late uninfected control female house flies. Bottom table: Early sporulation *E. muscae* female house flies compared to early uninfected control female house flies.

**Supplementary video 1**

https://sid.erda.dk/share_redirect/hIekF7iOhD

Timelapse video of sporulating female house fly. The video spans 24 hours.

To view video: Follow the link. This will download the videofile.

This file can be played with a media player (tested with VLC media player & Windows media player).

**Supplementary video 2**

https://sid.erda.dk/share_redirect/bJf0Bner0Q

Video of escaped house flies feeding off conidia on a petri dish lid and bottom. The conidia can be seen as a white powder like substance in the top of the lid.

To view video: Follow the link. This will download the videofile.

This file can be played with a media player (tested with VLC media player & Windows media player).

